# Cell adhesion and spreading on fluid membranes through microtubules-dependent mechanotransduction

**DOI:** 10.1101/2022.09.12.507658

**Authors:** Oleg Mikhajlov, Ram M. Adar, Maria Tătulea-Codrean, Anne-Sophie Macé, John Manzi, Fanny Tabarin, Aude Battistella, Fahima di Federico, Jean-François Joanny, Guy Tran van Nhieu, Patricia Bassereau

## Abstract

During cell adhesion, integrins form clusters that transmit mechanical forces to the substrate (mechanotransduction) and regulate biochemical signaling depending on substrate stiffness. Studies on mechanotransduction significantly advanced our understanding of cell adhesion and were mostly performed on rigid substrates. In contrast to rigid substrates, integrins’ ligands on fluid supported lipid bilayers (SLBs) are mobile and adhesive complexes cannot serve as anchoring points promoting cell spreading. Here, we demonstrate that cells spread on SLBs coated with Invasin, a high-affinity integrin ligand. We show that in contrast to SLBs functionalized with RGD peptides, integrin clusters grow in size and complexity on Invasin-SLBs to a similar extent as on glass. While actomyosin contraction dominates adhesion maturation on stiff substrates, we find that integrin mechanotransduction and cell spreading on fluid SLBs rely on dynein pulling forces along microtubules perpendicular to membranes and microtubules pushing on adhesive complexes, respectively. These forces that may also occur on non-deformable surfaces are revealed in fluid substrate set ups. Our findings, supported by a theoretical model, demonstrate a new mechanical role for microtubules in integrin clustering.

---

Integrin-mediated adhesion is critical to fundamental cellular processes such as cell migration^1^, differentiation^2^, and the development of tissues and organs^3^. Integrin clusters serve as communication hubs transmitting mechanical forces between cells and substrates. During integrin-mediated mechanotransduction, forces generated by cells and transmitted to a substrate regulate biochemical signaling as a function of substrate stiffness^4^.

In recent years, studies on mechanotransduction have brought a wealth of information and conceptual shifts in our understanding of cell adhesion^5^. A plethora of “adhesome proteins” connecting integrins with cytoskeleton were identified, and their role in mechanotransduction and cell adhesion was established^6^. The majority of these studies, however, were performed on substrates with immobilized integrin ligands, such as glass or deformable 2D-gels^7^. The actin cytoskeleton plays a crucial mechanical role in adhesion reinforcement on stiff substrates^8^. Actin polymerization is required to form small nascent adhesions (NAs)^9^, and actomyosin contraction promotes their growth into large and dense focal adhesions (FAs) connected to stress fibers^10^. However, much less is known about adhesion on soft substrates like 3D matrices^11^ or to the plasma membrane of other cells^12^, relevant to the interaction between immune and target cells^13,14^, where integrin ligands are not immobilized but embedded on fluid membranes.

Supported lipid bilayers (SLBs) are convenient model membranes for mimicking the fluid characteristic of plasma membranes to study cell-cell adhesion^15^. Previous studies with SLBs functionalized with canonical RGD peptides have shown that cells neither spread nor develop large and dense integrin adhesions like FAs and stress fibers^16,17^. The current understanding is that mobile integrin-ligand complexes on bilayers cannot serve as anchoring points promoting cell spreading as on stiff substrates^10^, and NAs cannot reinforce through mechanotransduction to promote strong adhesion on fluid substrates^18^. However, previous studies have not addressed the effect of the integrin receptor-ligand affinity that may significantly differ from RGD peptides and could regulate cell adhesion^19,20^.

To directly test the role of integrin receptor-ligand affinity, we used SLBs functionalized with a high-affinity ligand, the *Yersinia* bacterial protein Invasin^21^, that binds to a subset of β_1_-integrins, including the fibronectin receptor α_5_β_1_. On these SLBs, we seeded mouse embryonic fibroblasts (MEF) expressing a recombinant integrin β_1_-subunit labeled with a Halotag at its ectodomain instead of the endogenous β_1_-integrin^22^. This construct conjugated with membrane impermeable Halotag-dyes allows studying β_1_-integrins primarily at the cell surface, excluding signals from integrins inside the cell observed with genetic labeling. We used confocal microscopy to detect β_1_-integrin clustering during the time course of adhesion. We quantified cluster areas and integrin densities using fluorescent SLB calibration standards^15^ and compared them for Invasin- and RGD-SLBs. Accordingly, we provide evidence for mechanotransduction occurring on Invasin-SLBs, leading to FA-like β_1_-integrin adhesions. Mechanotransduction occurs on fluid SLBs because of vertical forces acting on integrin clusters generated by microtubules and dyneins. We have developed a theoretical model supporting this new mode of mechanotransduction. Additionally, we found that cells spread on fluid Invasin-but not on RGD-SLBs. Finally, we show that microtubules but not actin play a significant role in mechanotransduction on these SLBs.

## Results

### Cell “trembling” and spreading on SLBs depend on ligands

SLBs were functionalized with RGD peptides or Invasin, and their fluidity was confirmed by FRAP (Supplementary Fig. S1). Following the addition of MEF cells using brightfield microscopy, we observed cells with fluctuating edges that we called “trembling”, as well as “adherent” cells with immobile edges (Fig. 1A; Supplementary videos V1-V2). The fraction of the latter increased over time, reaching 80% on RGD- and Invasin-SLBs after 45 minutes (Fig. 1B). The fraction of “adherent” cells increased faster on RGD- than on Invasin-SLBs (Fig. 1B), possibly due to higher effective RGD densities on SLBs (20,000 RGD/µm^2^ vs. 600 Invasin/µm^2^) and/or to the RGD binding to a larger range of integrin types that could facilitate integrin activation^23^. To further confirm this, we studied cell adhesion dynamics in the presence of manganese (Mn^2+^) that activates integrins^24^. Strikingly, integrin activation by Mn^2+^ did not affect adhesion dynamics on RGD-SLBs. In contrast, it significantly accelerated adhesion on Invasin-SLBs, which had a similar kinetics as on RGD-SLBs (Fig. 1B). While similar cell edge fluctuations were previously reported on glass^25^, we found, however, that the adherent proportion on glass remained constant and similar for both ligands with and without Mn^2+^ (Supplementary Figs. S2A-C). These observations suggest that cell adhesion on SLBs but not glass depends on the cell ligand’s ability to activate integrins.

**Figure 1.**
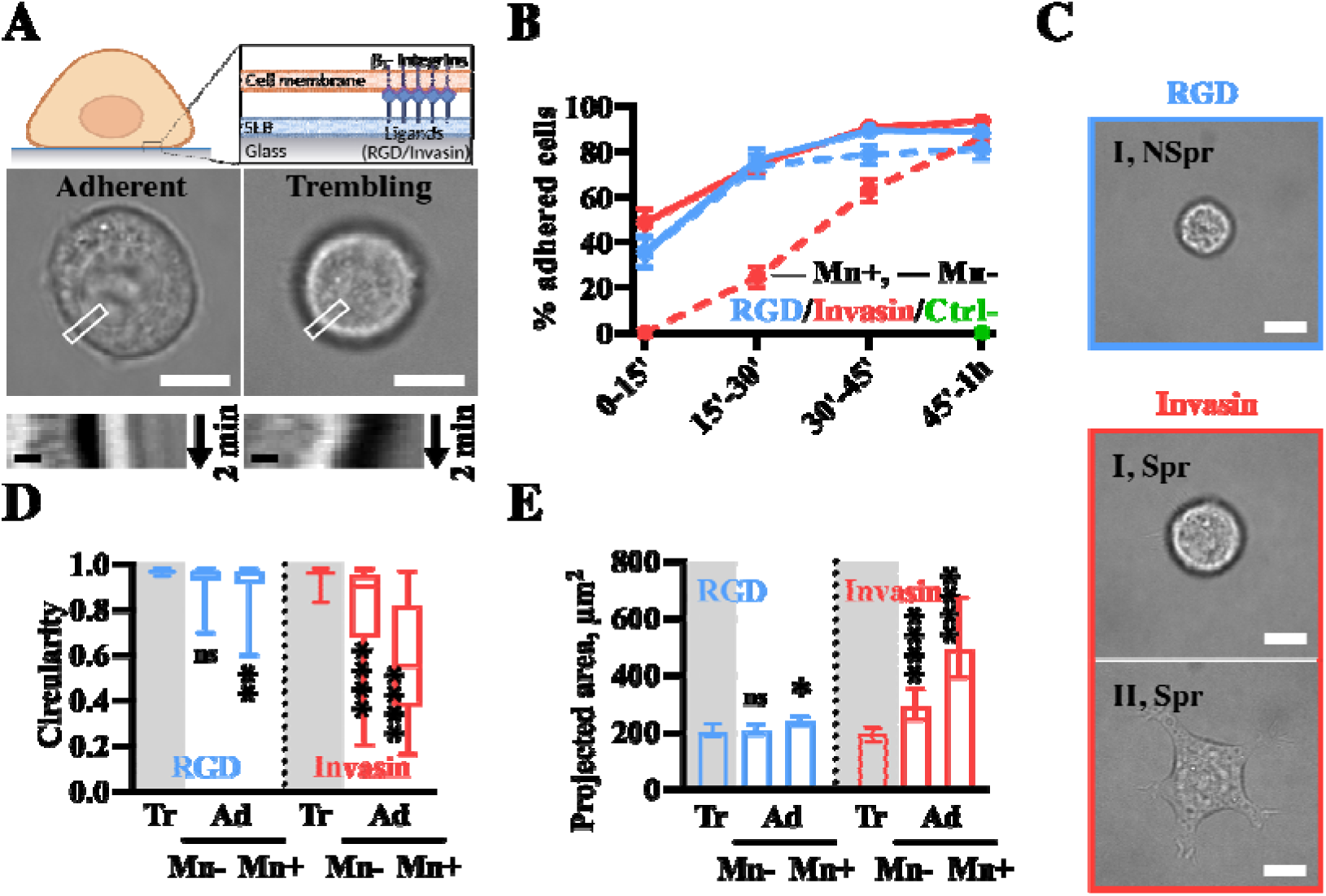
MEF cells adhere faster on RGD-SLBs but spread more on Invasin-SLBs. A) Top: schematic illustration of a cell adhering on an SLB coated with a ligand of β_1_-integrin (RGD peptide, or Invasin) (created with BioRender.com). Bottom: brightfield images of an adherent (left) and a trembling (right) cell on Invasin-SLB with corresponding kymographs describing cell edge movements over 2 minutes. Scale bars: 10 µm; 1 µm. B) Time evolution of fractions of adherent cells on SLBs functionalized with RGD (blue) and Invasin (red) in presence (Mn+, full line) or absence (Mn-, dashed line) of Mn^2+^. Each data point represents between 42 and 113 cells studied in at least 3 independent experiments. A fraction of adherent cells 45 minutes after seeding on SLBs without ligands (Ctrl-, green) in absence of Mn^2+^. Line scatter plot, mean, SEM. C) Brightfield images of cells with different morphologies (I – round (circularity > 0.8); II – not round (circularity < 0.8); Spr – spread (projected area > 200 µm^2^) and NSpr – non-spread (projected area < 200 µm^2^)) on SLBs functionalized with RGD (blue) and Invasin (red). Scale bars: 10 µm. D-E) Trembling (“Tr”, plots on grey background area) and adherent (“Ad”, 45 min – 1h after seeding) cells in presence (Mn+) or absence (Mn-) of Mn^2+^ adhering on RGD-(blue) or Invasin-coated (red) SLBs. Data from 44 cells, N=3 for trembling cells on RGD; 57 cells, N = 3 for trembling cells on Invasin (both Mn-); 49 cells, N=3 for cells adhering on RGD (Mn-); 53 cells, N=3 for cells adhering on Invasin (Mn-); 58 cells, N=3 for cells adhering on RGD (Mn+) and 49 cells, N=5 for cells adhering on Invasin (Mn+). D) Circularity of trembling and adherent cells, based on cell edge detection from brightfield images, 45 minutes after seeding. Box plot. E) Projected area of the cells described in D. Bar plot, median, 95% CI.

Next, we analyzed the morphology of cells spreading on SLBs as a proxy for cell adhesion strength^26^ by measuring their “projected areas” on the SLB surface and their “circularities”, which characterize cell shape irregularity (Fig. 1C). Consistent with an early adhesion stage, all “trembling” cells on both ligands had a relatively small and round projected area, independently of Mn^2+^ treatment (Supplementary Figs. S2D-E). Moreover, in agreement with previous studies on fluid substrates^16^, we found that fibroblasts did not spread on RGD-SLBs, keeping small projected areas (<200 µm^2^) and remaining round (0.8 < circularity < 1), independently of Mn^2+^ treatment (Figs. 1D-E). In contrast, cells spread significantly more on Invasin-SLBs, with median projected areas 1.5- and 2-fold higher than on RGD-SLBs for Mn^2+^-untreated and treated cells, respectively (Fig. 1E). Surprisingly, 35% of Mn^2+^-untreated and 75% of Mn^2+^-treated cells on Invasin-SLBs had irregular shapes with multiple protrusions (circularity < 0.8) and projected areas twice and 3-fold higher than the trembling reference, respectively (Fig. 1C).

Altogether these results indicate that while integrin activation might be a limiting factor on Invasin-SLBs, these can promote cell spreading upon integrin activation by Mn^2+^, in contrast to RGD-SLBs.

### β_1_-integrin clusters are denser and larger on Invasin-than on RGD-SLBs

We used confocal microscopy to study the clustering of β_1_-integrins labeled with Alexa488-Halotag ligands at the cell-SLB interface during the first hour of adhesion. For both RGD- and Invasin-SLBs, we observed the formation of β_1_-integrin clusters with isotropic shapes, morphologically different from FAs or actin-dependent podosome-like structures mainly composed of β_3_-integrins (Pr. Cheng-Han Yu, private communication)^17^. We applied fluorescence calibration to determine integrin density maps (Supplementary Fig. S3), and image segmentation programs to detect integrin clusters and measure their areas (σ) and densities of integrins (ρ) (Methods). Specifically, we segmented integrin density maps with a threshold of 300 integrins/µm^2^, corresponding to the minimal spacing of 58 nm between integrin-ligand pairs observed during mechanotransduction on glass^27^.

We observed that 45 minutes after cell seeding, clusters on Invasin-SLBs occupied larger areas and were denser than on RGD-SLBs independent of Mn^2+^ (Figs. 2B-D). Median densities of integrins per cell (ρ) reached about 100 integrins/µm^2^ for trembling cells for both ligands (Fig. 2B; Supplementary Figs. S4A-B). However, the median ρ had a significant higher increase for adherent cells than for trembling cells: up to 160 integrins/µm^2^ and 450 integrins/µm^2^ on RGD- and Invasin-SLBs, respectively. Noteworthy, ρ was significantly higher for Invasin-than RGD-SLBs, reaching values comparable to glass^28^, that were never observed for RGD-SLBs (Fig. 2B). Moreover, individual integrin clusters were more than twice denser on Invasin- than on RGD-SLBs, with mean values equal to 300 integrins/µm^2^ and 125 integrins/µm^2^ on Invasin and RGD, respectively (Fig. 2C). Additionally, we compared areas of dense integrin clusters σ_300_, corresponding to ρ > 300 integrins/µm^2^ (Fig. 2D). Independent of Mn^2+^ treatment, σ_300_ per cell was significantly higher on Invasin-SLBs, with 23% of clusters larger than the theoretical diffraction limit, DL (183 nm; Methods), as opposed to only 9% for RGD-SLBs (Fig. 2E).

**Figure 2.**
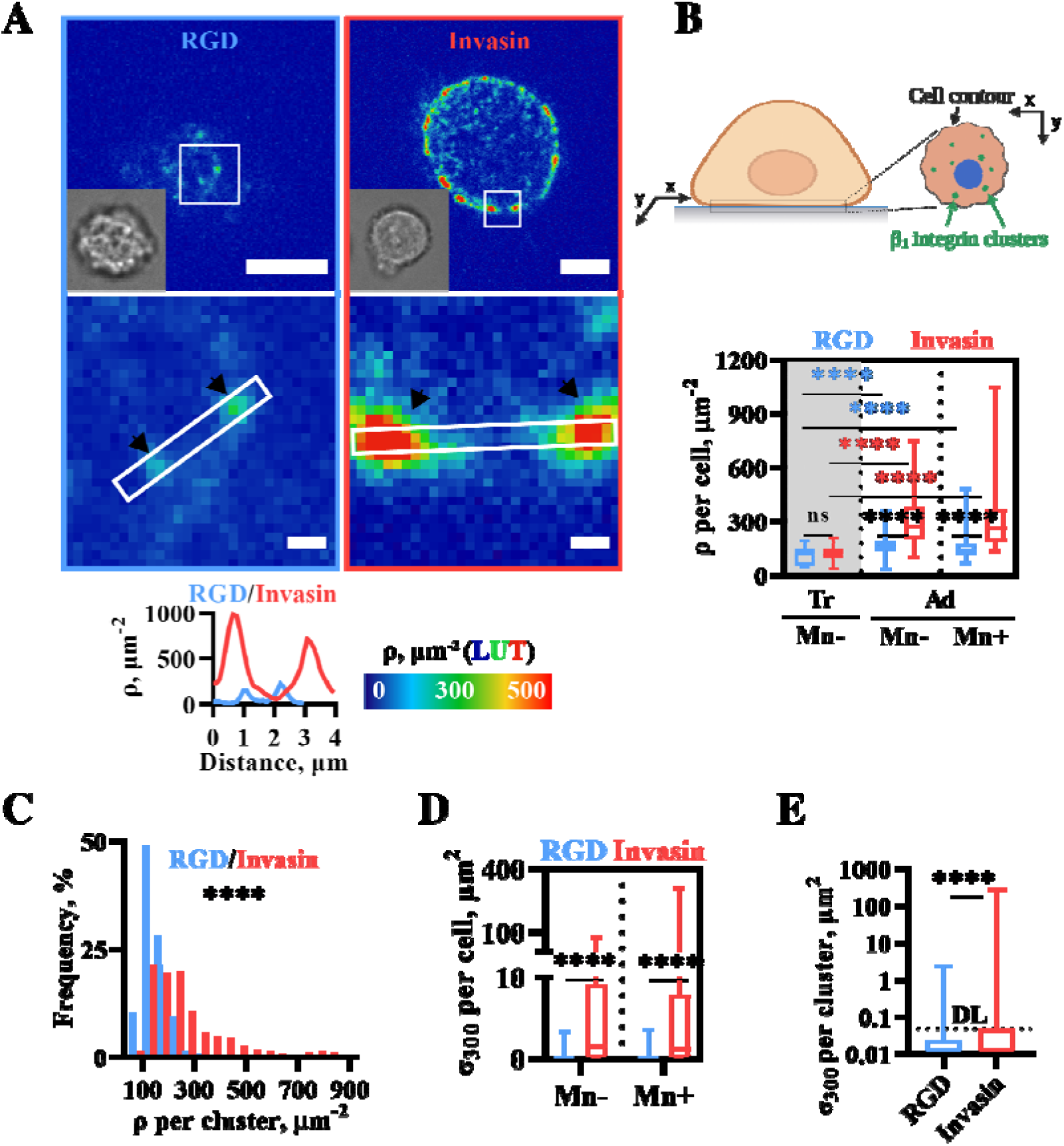
β_1_-integrin clusters in MEF cells adhering on SLBs and relationship between their density and cell spreading. A) Main panels: β_1_-integrin density maps, using the “physics” LUT of ImageJ below (cf. Methods for the calibration derived from fluorescence images), showing β_1_-integrin cluster organization in cells adhering on SLBs coated with RGD (upper left) and Invasin (upper right). Images are taken at the focal plane of the SLBs. Lower left corners: corresponding brightfield images. Zoomed panels (white squares on the main panels): regions with β_1_-integrin clusters (black arrows). Bottom: β_1_-integrin density profiles along the white rectangles in the zoomed panels. Scale bars: 5 µm (main panels); 1 µm (zoomed panels). B-E) RGD-SLB (blue), Invasin-SLB (red). Data from 57 cells, N=3 for trembling cells on Invasin; 44 cells, N=3 for trembling cells on RGD (both Mn-); data from adhering cells 45 min – 1h after seeding : 83 cells, N=4 for cells adhering on Invasin (Mn-); 48 cells, N=3 for cells adhering on RGD (Mn-); 78 cells, N=5 for cells adhering on Invasin (Mn+) and 102 cells, N=4 for cells adhering on RGD (Mn+). B) Top: schematic illustration of integrin clustering at the cell-SLB interface (created with BioRender.com). Bottom: Distributions of mean β_1_-integrin density per cell in trembling (“Tr”, plots on grey background area) and adherent (“Ad”) cells in presence (Mn+) or absence (Mn-) of Mn^2+^ adhering on RGD- or Invasin-coated SLBs. Box plot. C) Histograms of integrin densities in individual clusters in Mn^2+^ treated cells adhering on RGD- or Invasin-SLBs. Data from 11,441 clusters on Invasin and 16,115 clusters on RGD. D) Distributions of total area of β_1_-integrin clusters of density higher than 300 β_1_-integrins/µm^2^ per cell (σ_300_) in the presence (Mn+) or the absence (Mn-) of Mn^2+^ in cells adhering on RGD- or Invasin-SLBs. Box plot. E) Distributions of areas of individual β_1_-integrin clusters of density higher than 300 β_1_-integrins/µm^2^ in Mn^2+^-treated cells adhering on RGD (blue)- or Invasin (red)-coated SLBs. 75-percentile of σ_300_ per cell distribution: 8-9 µm^2^ for Invasin-SLBs and 0.1-0.15 µm^2^ for RGD-SLBs. DL – theoretical diffraction limit. Data from 347 clusters for cells adhering on Invasin and 8,954 clusters for cells adhering on RGD. Box plot.

### FA proteins are recruited at β_1_-integrin clusters

Integrin clusters grow and mature during cell adhesion on rigid substrates, recruiting FA proteins in response to mechanotransduction driven by actomyosin contractility^6,29^. To determine whether β_1_-integrin adhesive clusters could also mature on SLBs, we quantified the recruitment of FA proteins (Figs. 3A-B). “Early adhesion proteins” like talin, kindlin-2, paxillin, vinculin and “late adhesion proteins” like VASP and zyxin were recruited to β_1_-integrin clusters both on Invasin- and RGD-SLBs (Supplementary Figs. S4A-E). Except for talin, however, we observed a higher enrichment of all FA proteins on Invasin-than on RGD-SLBs, consistent with higher mechanical forces applied on Invasin-bound cells.

**Figure 3.**
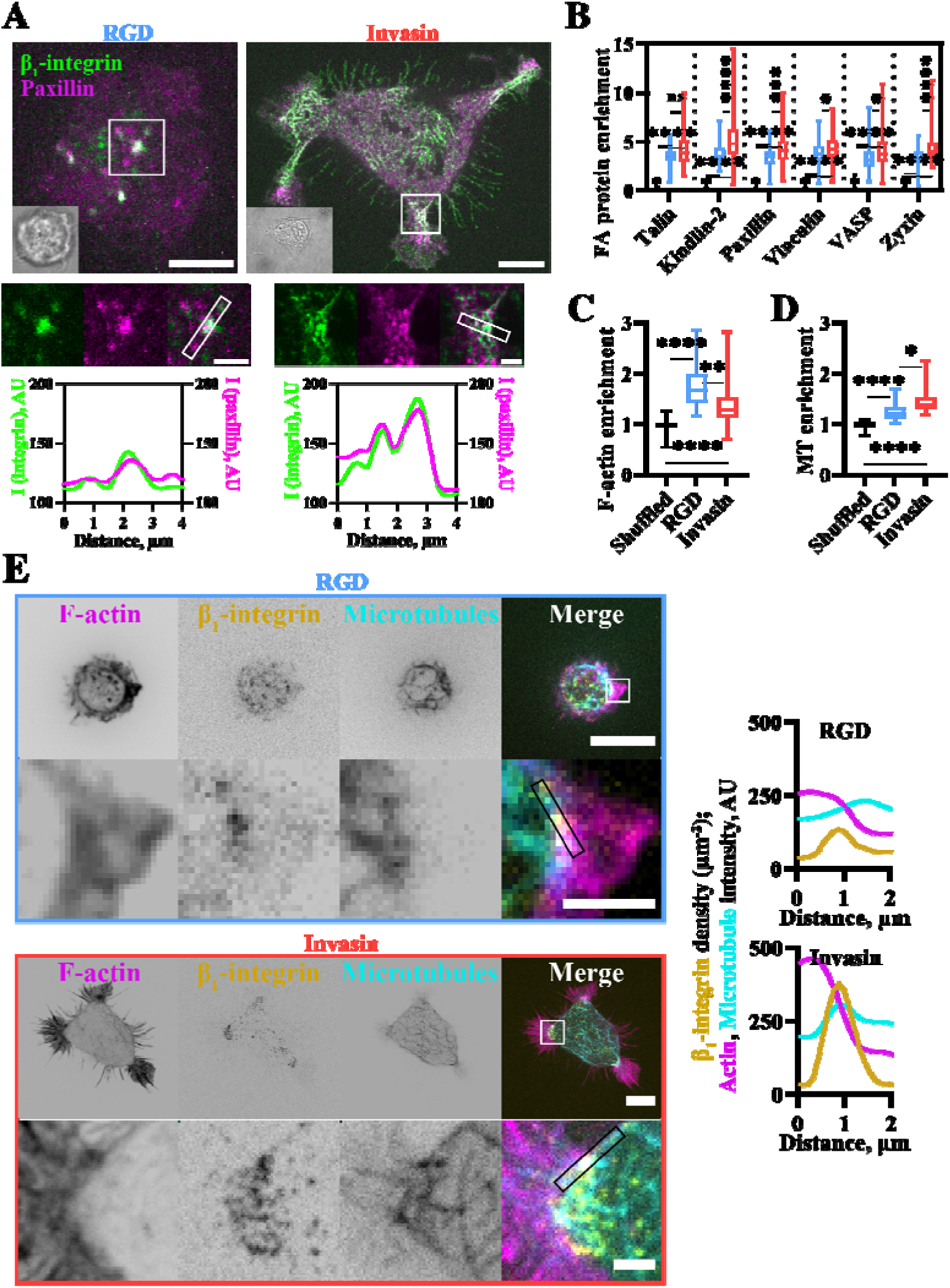
Focal adhesion proteins and cytoskeleton (actin and microtubules) are enriched at β_1_-integrin clusters on SLBs. All cells are Mn^2+^-treated and imaged 45 min – 1h after seeding in the chamber. A) Main panels: fluorescence multi-channel images showing β_1_-integrin (cyan) and paxillin (magenta) cluster organization in cells adhering on SLBs coated with RGD (blue frame, left) and Invasin (red frame, right). Images are taken as max intensity z projections (in Fiji) of the volume with 1 µm height (z coordinate) centered at the focal plane of the SLB. Upper left corners: corresponding brightfield images. Medium panels: zooms corresponding to the white squares on the main panels. Regions with β_1_-integrin and paxillin clusters. Bottom: β_1_-integrin and paxillin intensity profiles along the lines (white rectangles in the zoomed panels). Scale bars: 5 µm (main panels); 1 µm (zoomed panels). B) Enrichment of FA proteins in β_1_-integrin clusters in cells adhering on SLBs functionalized with RGD (blue) and Invasin (red) as a mean fluorescence intensity of the FA protein in the region of β_1_-integrin clusters divided by the mean fluorescence intensity of the protein in the cell. It was compared with “shuffled” control enrichments (black), enrichment calculated at random pixels instead of β_1_-integrin clusters regions. Box plot. Data for every FA protein from between 108 and 181 cells for the shuffled control (shuffled), between 42 and 87 cells (RGD) and between 56 and 128 cells (Invasin). All results are representative of at least 3 independent experiments. C) Enrichment of F-actin in β_1_-integrin clusters in cells adhering on SLBs functionalized with RGD and Invasin. Box plot. Data from 143 cells (shuffled), 58 cells (RGD) and 85 cells (Invasin). N ≥ 3. D) Enrichment of microtubule in β_1_-integrin clusters in cells adhering on SLBs functionalized with RGD (blue) and Invasin (red). Box plot. Data from 107 cells (shuffled), 53 cells (RGD) and 54 cells (Invasin). N ≥ 3. E) Fluorescence multi-channel images showing β_1_-integrin (yellow), actin (magenta) and microtubules (cyan) in cells adhering on SLBs coated with RGD (blue frame, left) and Invasin (red frame, right). Images are taken in the focal plane of the SLB. β_1_-integrin density, actin and microtubules intensity profiles along the lines (black rectangles in the zoomed panels) are plotted for both cells. Scale bars: 5 µm, 1 µm.

While integrin-mediated mechanotransduction on glass is generally associated with the actin cytoskeleton, microtubules can stabilize and regulate protein turnover in FAs^30,31^. We first measured the association of F-actin to β_1_-integrin clusters in MEF cells using lifeact-mScarlet (Fig. 3C). We found actin enrichment at β_1_-integrin clusters for both ligands but did not observe actin stress fibers usually associated with mechanotransduction (Figs. 3C; 3E). Furthermore, we found that F-actin was more enriched at integrin adhesion clusters on RGD-than Invasin-SLBs (Fig. 3C). These results contrast with the difference in the enrichment of FA proteins at these structures and suggest that actin does not drive FA maturation on Invasin-SLBs. We then investigated the role of the microtubule cytoskeleton using EMTB-iRFP. We found that microtubules were organized in similar networks at the adhesion interface on SLBs and glass (Supplementary Figs. S5A-B). However, microtubules were enriched at β_1_-integrin clusters with a 17% higher median on Invasin-compared to RGD-SLBs (Fig. 3D), suggesting their involvement in maturation. Moreover, we observed that adaptor proteins that connect talin to microtubules (Kank1^32^ and ELKS^33^) were recruited to β_1_-integrin clusters (Supplementary Figs. S5C-D).

### Dynein pulling along vertical microtubules leads to growth of integrin clusters

During mechanotransduction, integrin clusters are subjected to cellular mechanical forces that significantly increase in magnitude if applied against rigid substrates^10^. On fluid SLBs, lateral components of the forces are negligible compared to glass and are only associated with membrane viscosity^34^. However, SLBs can resist higher magnitude normal forces leading to the maturation of integrin clusters. These normal forces might lead to coupled local deformations of SLBs and cell plasma membranes. We observed such deformations while imaging β_1_-integrin clusters above the SLB focal plane (Fig. 4A; zoom 1; 0 < z < 1.5 µm). Akin to clusters at SLB surfaces (Fig. 4A; zoom 2; z = 0), clusters in the cell volume had a larger total area in cells on Invasin-than on RGD-SLBs (Fig. 4B). Some of them were associated with membrane tubes pulled out of SLBs and locally vertical actin and microtubules associated with them (Fig. 4A; zoom 2). The proportion of cells with tubes and the number of tubes per cell were significantly higher on Invasin than on RGD (Fig. 4C). These results are consistent with the notion that integrin clusters on Invasin-SLBs are exposed to higher mechanical forces than those on RGD-SLBs, leading to their growth in size and density, higher recruitment of FA proteins and more tubes.

**Figure 4.**
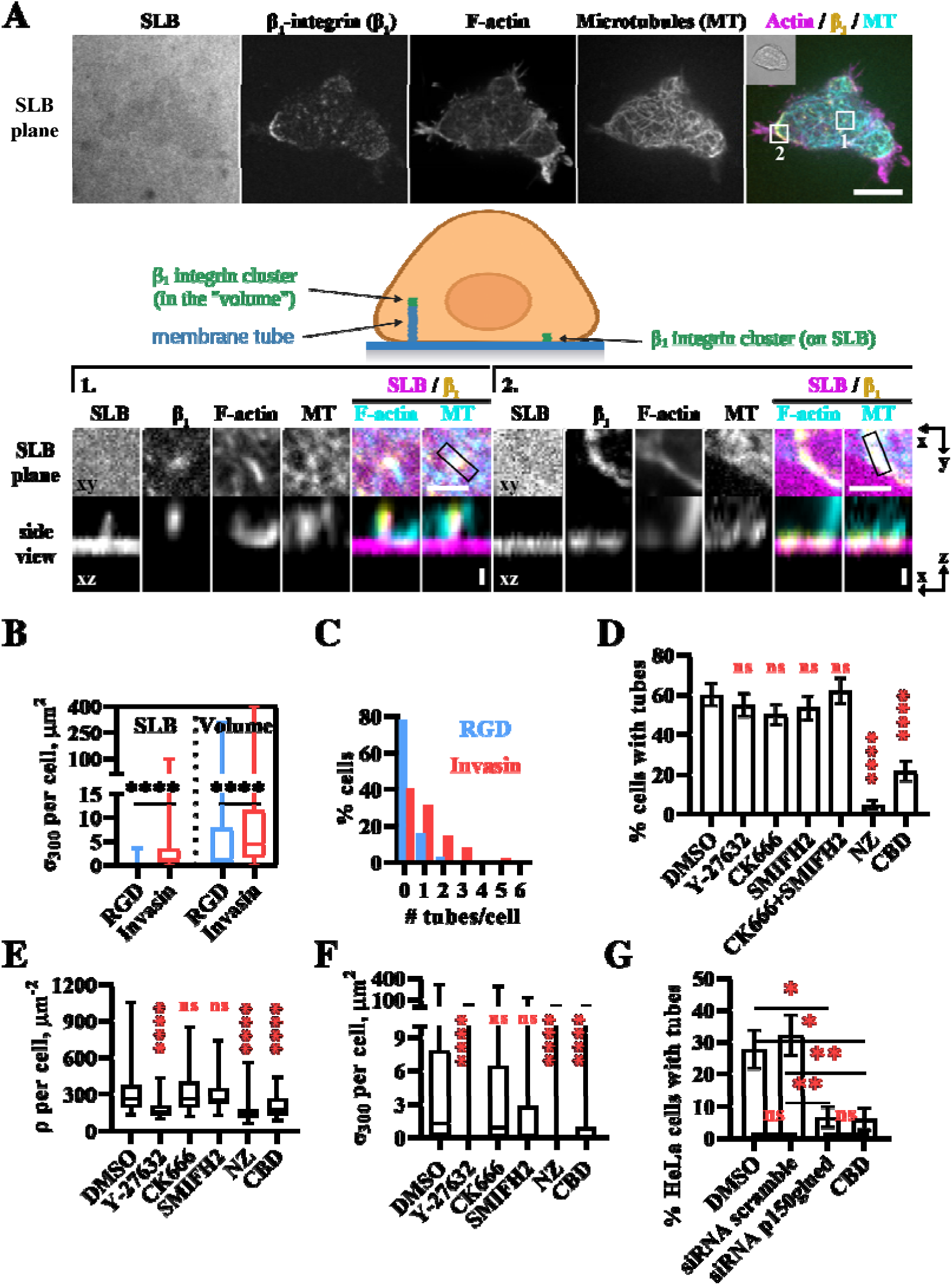
Tube formation at β_1_-integrin clusters and cluster densification is driven by microtubules and dynein activity. All cells are Mn^2+^-treated and imaged 45 min – 1h after seeding in the chamber on Invasin-SLBs. A) Top panel: fluorescence multi-channel image of a MEF cell showing the SLB labeled with a TR-DHPE lipid, β_1_-integrin with Halotag-Alexa488, actin with lifeact-mScarlet and microtubules (MT) with EMTB-iRFP. A brightfield image of the cell is in the upper left corner of the merged image. Middle panels: schematic illustrations of a cell with integrin clusters associated with a membrane tube pulled from the SLB (left) and resided on the surface of the SLB (right) (created with BioRender.com). Bottom panels (zoom): regions corresponding to white squares 1 and 2 on the main panels. Top: x-y. Bottom: x-z section corresponding the lines in x-y. Scale bars: 10 µm (main panel); 2 µm (zoomed panel xy); 0.5 µm (zoomed panel xz). B-C) Invasin-SLBs: 78 cells, N=4; RGD-SLBs: 102 cells, N=4. B) Distributions of total area of β_1_-integrin clusters per cell at the SLB level and in the volume 1.5 µm above the SLB for MEF cells adhering on RGD- (blue) and Invasin-coated (red) SLBs. Box plots. C) Histograms of the number of detected tubes per cell in MEF cells adhering on RGD- (blue) and Invasin-coated (red) SLBs. D) Proportion of MEF cells with tubes. Cells adhere on Invasin-coated SLBs in presence of drugs. Data from 78 cells, N=4 for DMSO; 104 cells, N=4 for CK666; 71 cells, N=3 for SMIFH2; 58 cells, N=3 for CK666+SMIFH2; 72 cells, N = 3 for Y27632; 86 cells, N=3 for Nocodazole (NZ); 69 cells, N=4 for Ciliobrevin D (CBD). Bar plot: mean, SEM. E-F) MEF cells treated with Nocodazole (NZ), Cilibrevin D (CBD) and non-treated (DMSO). Data from 86 cells, N=3 for Nocodazole (NZ); 69 cells, N=4 for Ciliobrevin D (CBD); 78 cells, N=4 for DMSO. E) Distributions of the mean integrin density per cell. Box plot. F) Distributions of the total area of β_1_-integrin clusters per cell. Box plot. G) Proportion of HeLa cells with tubes. HeLa cells were treated with siRNA against p150glued, siRNA scramble (as a negative control) and with CBD. Bar plot: mean, SEM. Data from 58 cells, N=3 for DMSO; 56 cells, N=3 for siRNA scramble; 61 cells, N=4 for siRNA p150glued and 50 cells, N=4 for CBD.

To characterize the origin of vertical forces pulling on integrin clusters, we tested the effects of inhibitors of cytoskeleton polymerization and associated motors on the proportion of cells with tubes and the number of tubes per cell on Invasin-SLBs that have a higher number of tubes compared to cells on RGD-SLBs (Fig. 4D; Supplementary Fig. S6A).

We found that the inhibition of formins and of the Arp2/3 complex that nucleate actin polymerization using SMIFH2 and CK666, respectively, had no effect on tube formation frequency, neither on integrin clustering (Fig. 4D-F; Supplementary Fig. S6A). Preventing actomyosin contractility by Rho kinase inhibition (Y-27632) or blebbistatin did not influence the frequency of tube formation in agreement with previous studies on RGD-SLBs^35^ (Fig. 4D). Y-27632, however, led to a significant decrease in β_1_-integrin cluster size and density (Figs. 4E-F).

Together, these results suggest that the forces associated with actin polymerization and actomyosin contractility are not critical for vertical deformations of the membrane. However, the effects of Y-27632 suggest that actomyosin play a role through another mechanism of mechanotransduction, perhaps related to the described local “pinching” of nascent adhesion clusters^36^.

Next, we investigated whether microtubules and associated molecular motors are involved in tubular deformations of SLBs, as their role in mechanosensing of FAs on rigid substrates was reported^37,38^. We observed dense and large integrin clusters at the SLB surface and on tubes associated with microtubules (Figs. 3D; 4A; Supplementary Fig. S5B). When microtubules were depolymerized with nocodazole (NZ) or when dynein activity was blocked with ciliobrevin D (CBD), the frequency and number of tubes decreased drastically (Fig. 4D; Supplementary Fig. S6A). Similar observations were made in HeLa cells on Invasin-SLBs after dynein inhibition with CBD or silencing of the dynactin subunit p150glued (Fig. 4G; Supplementary Figs. S6B-D). These findings suggest that vertical forces applied to β_1_-integrin clusters depend on microtubules and are driven by dynein activity. Finally, NZ and CBD treatment significantly decreased the total area of β_1_-integrin clusters and density (Figs. 4E-F). Noteworthy, this NZ effect is opposite to that observed on rigid substrates, where it leads to an increase in size and density of FAs due to enhanced actomyosin contractility ^39,40^.

Globally, these findings suggest that mechanotransduction on SLBs results from the pulling activity of the dynein motors on locally vertical microtubules that can resist mechanical forces (Supplementary Fig. S5B), leading to the growth and maturation of integrin clusters. Higher microtubule enrichment and higher frequency of tube formation on Invasin- than on RGD-SLBs suggests that mechanotransduction at integrin clusters on SLBs is affinity-dependent, in contrast with glass. This is consistent with longer lifetime ligand-integrin bonds supporting more efficient mechanotransduction.

To further characterize the role of integrin-ligand affinity on integrin clustering on fluid substrates, we propose a theoretical model (Supplementary Material) that supports the fact that ligand-receptor pairs with higher affinity form clusters of larger density and can sustain larger vertical forces (Supplementary Fig. S7). The model considers the concentrations of ligands, receptors, and bonds (i.e., bound ligand-receptor complexes) and the average distance between the cell membrane and SLBs. SLB fluidity allows for ligand (and possibly bond) mobility, which plays a dual role in the theory. First, ligand diffusion acts as a chemical contact with a reservoir and promotes cluster formation. Second, the large tangential forces exerted by the cytoskeleton tend to displace bonds rather than break them. Hence, the vertical cytoskeletal forces are the primary driver of mechanotransduction in our theory. We consider adhesion clusters as dense phase-separated regions of bonds. Phase separation is driven by the interplay between diffusion and attractive bond-bond interactions via two mechanisms. The first attractive mechanism is mediated by the membranes^41–43^ and the second by adaptor proteins in the adhesion site^44–46^. Cytoskeletal forces drive conformational changes in adaptor proteins and increase the effective attraction between bonds (e.g., via the exposure of vinculin binding sites on talin^47,48^). The formation of adhesion sites depends on affinity and vertical cytoskeletal force in the following way: forces increase the bond-bond attraction, and affinity increases the total number of bonds (Supplementary Fig. S7). The system is predicted to phase separate for sufficiently numerous bonds and strong attraction, implying adhesion clusters. The phase diagram obtained from the model agrees with our experimental results of cell adhesion on SLBs, for which the model explains the differences in clustering between the two ligand types (Supplementary Fig. S7).

### Integrin clusters are pushed to the cell periphery in a dynein- and microtubule-dependent manner

Cell spreading correlates with integrin clustering due to mechanotransduction on glass and high viscosity RGD-SLBs^34^. Unexpectedly, we observed that cells formed protrusions on fluid Invasin-SLBs, leading to their symmetrical spreading without polarization (Fig. 5A). We found that high integrin densities in clusters in Mn^2+^-treated cells on Invasin-SLBs correlated with high spreading area and low cell circularity (Figs. 5B; Supplementary Fig. S8). Moreover, we measured and compared cluster areas at the cell periphery and center (two zones of equal areas and separated by a line equidistant from the cell edge (Fig. 5C)); we found that integrin and paxillin clusters in spread (projected area > 450 μm^2^) and not round (circularity < 0.8) cells were more at the cell periphery than at the cell center as opposed to the clusters in non-spread and round cells (Fig. 5D; Supplementary Fig. S9A-B). Additionally, cells with a larger area fraction of clusters at the periphery than in the center were more spread and less round (Supplementary Fig. S9C-D). On average integrin clusters were closer to the cell border in cells with dense than sparse integrin clusters (Fig. 5E). These results suggest a relation between the growth of adhesion clusters, their movement to the cell periphery and cell spreading.

**Figure 5.**
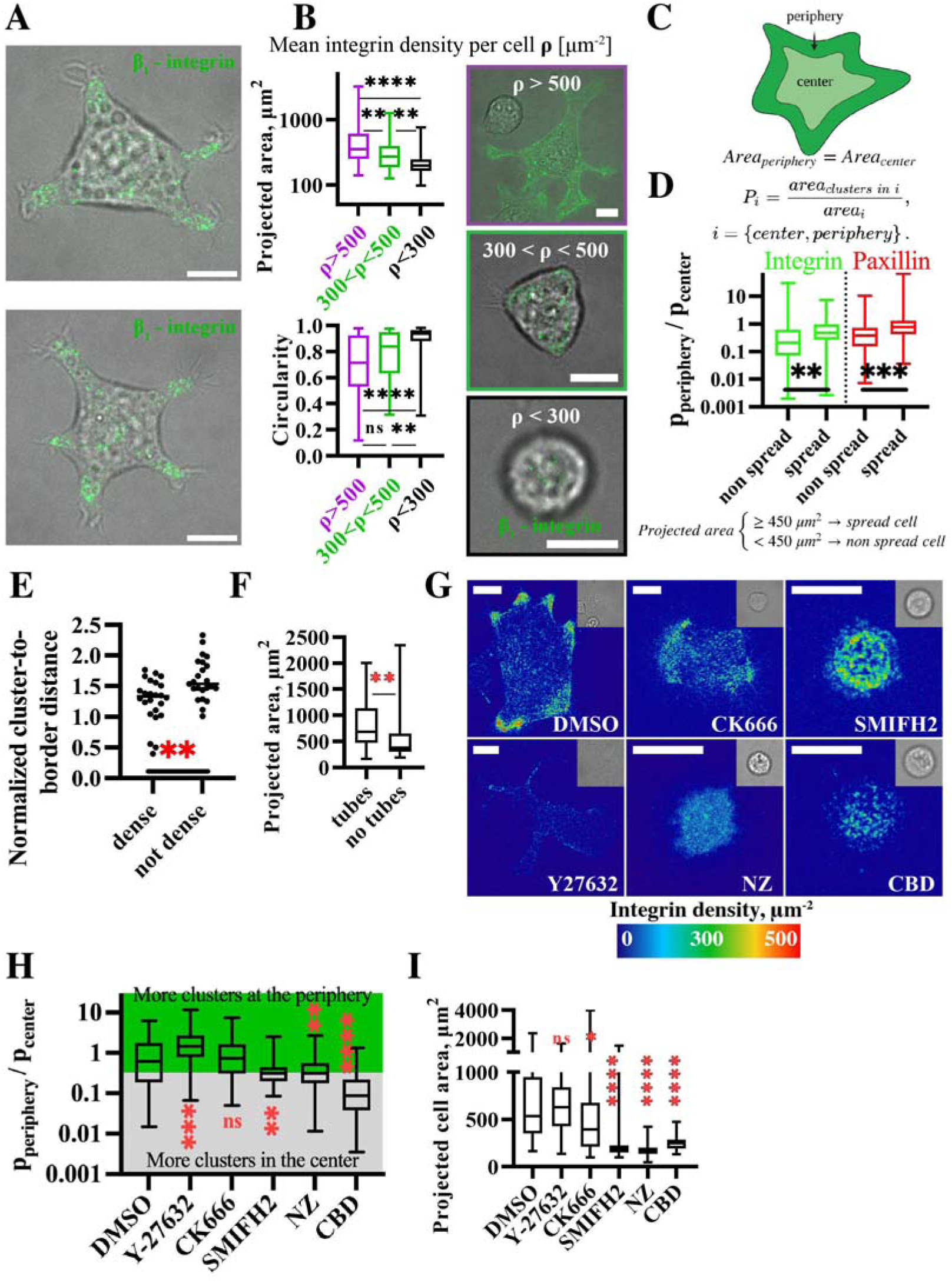
Dyneins push integrin clusters along microtubules to the cell periphery resulting in cell spreading on SLB. All cells are Mn^2+^-treated MEF cells and imaged 45 min – 1h after seeding to the chamber with Invasin-coated SLBs. A) Composite (brightfield and fluorescence channels) images of spread cells on Invasin-SLBs showing β_1_-integrin clusters (green). Scale bars: 10 µm. B) Projected cell area (top left) and cell circularity (bottom left) box plots of MEF cells with clusters of highest ρ > 500 (magenta); 300 < ρ < 500 (green); and ρ < 300 β_1_-integrins/µm^2^ (black). (right): typical composite (brightfield and fluorescence channels) images of spread cells on each of the three density classes showing β_1_-integrin clusters (green). Scale bars: 10 µm. Data from 157 cells studied in at least 3 independent biological experiments (N= 3) for ρ >500; 61 cells studied in at least N=3 for 300 <ρ < 500 and 87 cells studied in at least N=3 for ρ < 300. C) A schematic representation of the central and peripheral zones in a spreading cell (created with BioRender.com). Dark green: cell periphery that corresponds to the band parallel to the cell border; light green: cell center that corresponds to the part of the cell that excludes the cell periphery. The two zones are chosen such that they have the same area. D) Ratios of area fractions of β_1_-integrin (green) and paxillin (red) clusters in the periphery zone (P_periphery_) to the center zone (P_center_) for spread (projected area > 450 µm^2^) and non-spread (projected area < 450 µm^2^) cells. Box plots. (β_1_-integrin): N =4, 78 cells. (Paxillin): N=3, 119 cells. Mann-Whitney statistical test. E) Average distance between integrin clusters and the cell border normalized by average cluster-to-border distance for uniform integrin distribution for cells with high (ρ > 500 integrins/µm^2^) and low (ρ < 300 integrins/µm^2^) integrin density (Methods). A value above (or below) 1 describes an integrin cluster distribution skewed towards the center (or periphery) of the cell, relative to the uniform distribution. Scatter dot plot, median. Data from 24 cells, N=3 for cells with high integrin density, and from 23 cells, N=3 for cells with low integrin density. F) Projected cell area (µm^2^) of cells with and without detected tubes. Box plot. Data from 78 cells, N=4. G) Comparative illustration of integrin cluster distribution for the cells treated with drugs (CK666, SMIFH2, Y27632, NZ, CBD and DMSO for non-treated cells). β_1_-integrin density maps (represented with “physics” LUT of Fiji, calibrated bar on the side). Images are taken at the SLBs’ plane. Corresponding brightfield images are in the upper right corners. Scale bars: 10 µm. H-I) Data from 78 cells, N=4 with DMSO; 104 cells, N=4 with CK666; 71 cells, N=3 with SMIFH2; 58 cells, N=3 with CK666+SMIFH2; 72 cells, N=3 with Y27632; 86 cells, N=3 with Nocodazole (NZ); 69 cells, N=4 with Ciliobrevin D (CBD). H) Ratio of area fractions P of β_1_-integrin clusters in the periphery zone (P_periphery_) to the center zone (P_center_). Cells with more clusters in the periphery have P>1 and are in the green part of the graph where those with more clusters in the center have P<1 and are in the grey part. Box plot. I) Projected area of cells treated with drugs. Box plot.

Next, we found a positive correlation between cell spreading and the presence of membrane tubes, consistent with mechanotransduction at integrin clusters on SLBs (Fig. 5F). Moreover, spread cells had more tubes in the center than at the periphery (Supplementary Fig. S9E) – a result that indicated that more clusters in the center were subjected to vertical pulling forces than at the periphery, which agrees with recent findings of Brockman et al. on RGD-coated SLBs that integrin clusters experience more vertical forces in the cell center and more tangential forces at the cell periphery^49^. Because microtubules and dyneins were crucial for integrin cluster growth and densification, as well as the membrane tube formation on SLBs (Figs. 4E-F), we hypothesized that when oriented parallel to the substrate, they also play a role in the localization of clusters at the cell periphery, in a similar manner as for cells spreading on glass^50,51^. In line with this, we consistently observed accumulation of microtubules at peripheral integrin clusters in spread cells on Invasin-SLBs (Fig. 3E (Invasin); Fig. 4A (zoom 2); Supplementary Fig. S5B). Cell treatment with NZ or CBD decreased integrin clustering (Figs. 4E-F) and completely abolished integrin cluster localization at the cell periphery (Figs. 5G-H). Noteworthy, cells treated with NZ or CBD did not spread (Fig. 5I; Supplementary Fig. S9F). We interpret this result as a combination of inhibition of microtubule and dynein pushing on adhesion clusters and an increase in actomyosin contractility possibly linked to microtubule depolymerization^40^. Consistent with an antagonistic role of the actin cytoskeleton on cell spreading, inhibition of Rho kinase with Y27632 led to even more spread cells^52^ (Figs. 5G-I; Supplementary Fig. S9F). Similarly to cells on glass^53^ and RGD-SLBs^16^, inhibition of formins by SMIFH2 impaired cell spreading on Invasin-SLBs and integrin cluster movement to the cell periphery without significant effects on cluster growth (Figs. 4E-F; 5G-I; Supplementary Fig. S9F). These findings suggest that, due to dynein motor activity along microtubules, large and mature β_1_-integrin clusters are pushed towards the cell periphery, contributing to cell spreading on SLBs through actin-rich protrusions emanating from these adhesive clusters.

## Discussion

Our understanding of FAs is primarily based on studies on rigid substrates such as glass, where mechanotransduction is mainly driven by actomyosin-dependent forces transduced by actin fibers tangential to the cell basal membrane^54^. Living tissues, however, cover a wide range of stiffness^5^ and, instead of being immobilized, integrin ligands can be present in cell membranes^55,56^ where they may diffuse laterally with little constraints and may not exert the range of counter-forces required for “canonical” mechanotransduction. Indeed, when cells adhere on SLBs, the absence of actin stress fibers usually associated with FAs that planar forces on SLBs are negligible compared to glass^57,58^.

Using *Yersinia* Invasin as a high-affinity β_1_-integrin ligand, we show for the first time that cells adhere, spread, and develop large and dense β_1_-integrin clusters on fluid substrates (SLBs), similar to FAs on glass (Fig. 2B)^28^. Figure 6 summarizes our observations of cell adhesion on fluid substrates. Contrary to glass, on SLBs, actin-related forces tangential to the plasma membrane do not play a dominant role in integrin adhesion due to substrate fluidity. In addition, instead of enhancing adhesion and spreading through stimulation of actomyosin activity (observed on glass)^39^, microtubule depolymerization strongly inhibits it. This is very reminiscent of cells migrating on soft 3D extracellular matrix^50,59^. Microtubules were shown to be essential for cell spreading on relaxed collagen networks and for forming dendritic extensions^60^. We show here that adhesion maturation on bilayers relies on microtubule-dependent forces perpendicular to the bilayer. We evidence these forces by observing local SLB deformations/tubes connecting the bilayer and integrin clusters in the plasma membrane (Fig. 4A). The existence of normal force components at integrin clusters was also reported on RGD-SLBs using DNA-FRET force probes^49^, but their origin was not identified.

**Figure 6.**
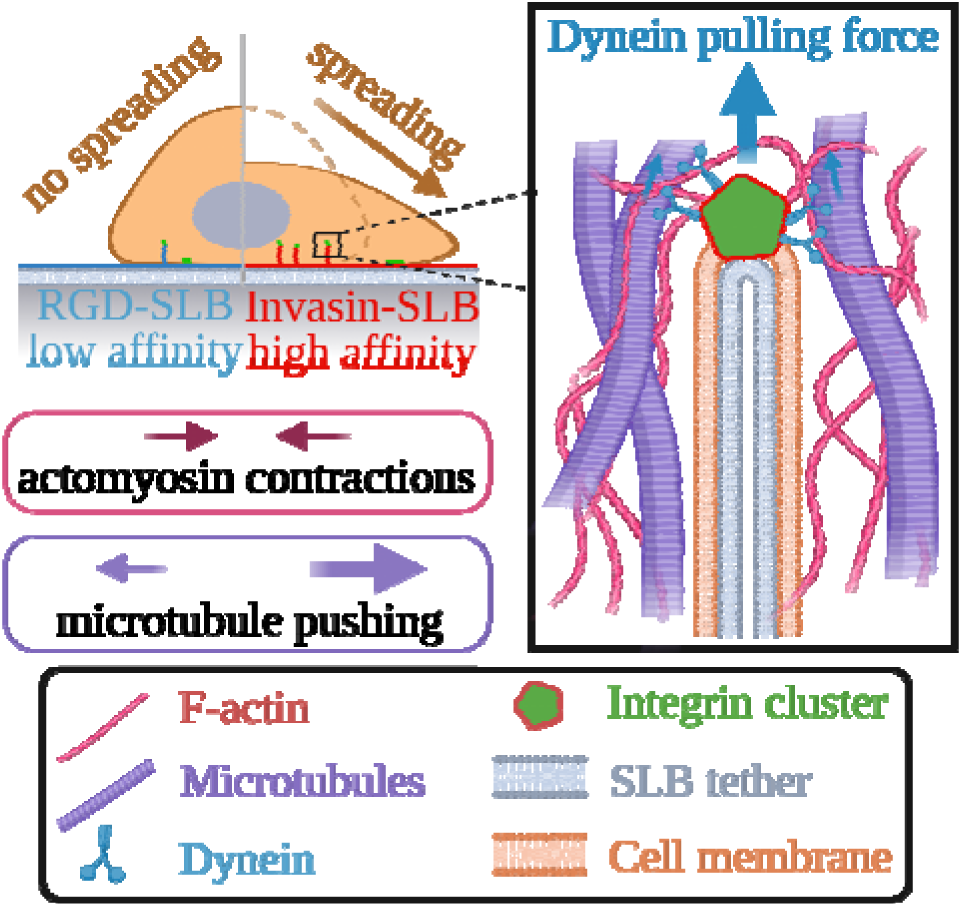
Role of receptor-ligand affinity in cell spreading and mechanotransduction during adhesion on SLBs. Created with BioRender.com

On glass, microtubules orient vertically^61^ and associate with mature FA at the cell periphery via several adaptor proteins^33,37^. Moreover, dyneins that stabilize microtubule plus-ends at the plasma membrane^62^ interact with integrin adhesion clusters through paxillin and other FA proteins^63,64^. Here, we present several pieces of evidence supporting a role for microtubules in exerting forces on β_1_-integrin clusters, normal to the SLB surface: i, microtubules are enriched at integrin clusters on SLBs (Fig. 3D); ii, microtubule-talin adaptors KANK1 and ELKS are recruited at these clusters (Supplementary Figs. S5D-E); iii, some microtubules have a vertical orientation in association with integrin adhesions (Fig. 4A).

Molecular motors can collectively pull tubes from membranes by distributing the load on a dynamically formed motor assembly, as shown *in vitro* for kinesins^65^. Forces of the order of 60 pN or less are necessary to pull tubes from SLBs on glass^66^. Dimers of dyneins exert forces of the order of 7 pN^67^ and can then pull on integrin adhesion complexes. When enough motors accumulate at the extremity of a microtubule, they produce the force required to tubulate together the plasma membrane and the SLB attached to it through integrin-ligand links. This force induces conformational changes in adaptor proteins leading to integrin clustering, which according to our theoretical model, is more efficient for high-affinity Invasin than RGD.

The observed local tubular membrane deformations connected to the clusters are microtubule- and dynein-dependent (but not actin-dependent) (Fig. 4D) and are not only involved in the cluster maturation but could also be related to β_1_-integrin endocytosis^68^. Active β_1_-integrin internalization proceeds via the clathrin-independent CLIC-GEEC (CG) pathway^68^. Interestingly, early stage of adhesion involves nanoclustering of GPI-anchored proteins (GPI-APs) as well as vinculin activation for a further maturation^69^. GPI-APs are also endocytosed though the same CG pathway^70^. These micrometer-long tubes could thus be non-cleaved endocytic structures. Indeed, contrary to glass, a double membrane tube is formed during endocytosis on SLBs, with the SLB membrane in the inner layer surrounded by the cell plasma membrane as an outer layer (Figs. 4A; 6). Such double-layer tubes could resist more to scission (dynamin-independent in the case of active β_1_-integrin^68^), leading to the formation of stable tubes.

On rigid substrates, cells form protrusions to spread. In this process, mature FAs act as anchoring points against which growing actin-rich protrusions push. The ability of cells to spread on glass depends on the density of ligands on the surface, the size of integrin adhesions, and their connection to the cytoskeleton^71^. On SLBs, on the other hand, integrin adhesions can move in the adhesion plane upon mechanical stimuli. On Invasin-SLBs, integrin clusters are dragged in the SLB plane towards the cell periphery in a microtubule and dynein-dependent manner (Fig. 3E). Since dyneins are involved in the maturation of the adhesions, essential for spreading, we cannot distinguish whether some dyneins also push microtubules linked to adhesions parallel to the surface towards the edge^72^ or whether microtubule dynamics is responsible for this. This movement is likely antagonized by an inward force due to the actomyosin contraction of the cell cortex^73,74^ (Figs. 3E; 5H; 6), leading to stalling of integrin clusters at the cell edge where they serve as anchoring centers from which actin-rich protrusions emanate^60,75^. The antagonistic effect of actin and microtubules may also be important in the densification of integrin clusters at the cell periphery, as we have found for well-spread cells (Figs. 5B-D). Importantly, such dense clusters are not found on Y-27632-treated cells (Figs. 4E-F; 5G). Cell protrusions on SLBs might result from a combination of pushing forces from actin and microtubule polymerization and microtubule pushing through dynein activity. The pushing forces are balanced against the centrosome to which microtubules are connected (Supplementary Fig. S5B). Remarkably, in contrast with cells on glass where microtubules are essential for cell polarization^76,77^, cells do not polarize on SLBs and conserve their central symmetry despite the central role of microtubules through a mechanism that remains to be characterized.

To summarize, both microtubules and actin play a role in the maturation of integrin clusters on SLBs. Microtubules and dynein motors exert vertical forces on integrin clusters, which result in the formation of tubes and are partially responsible for integrin clustering. At the same time, microtubules and dyneins exert lateral forces that push integrin clusters to the cell periphery. Cell spreading is determined by the interplay between the microtubule-driven pushing of integrin clusters and the actomyosin contractility at the cell edge. This interplay may also be responsible for the densification of integrin clusters at the cell periphery, as we found for well-spread cells.

In HeLa cells on SLBs, we also observed dynein/microtubule-dependent formation of tubes (Fig. 4G; Supplementary Figs. S6B-D), suggesting that mechanotransduction associated with forces normal to the adhesion plane occurs in various cell types. Functionalized SLBs allow to study the early stages of cell adhesion on a fluid interface, which is relevant for cell-cell interactions (i.e., brain cells, immune cells, etc.)^4^, but not accessible on substrates like glass or gels. Moreover, functionalized SLBs reveal forces probably at play on stiff substrates that cannot be observed with immobilized ligands. Using this platform, we found a new role for dynein motors and microtubules in integrin adhesion cluster growth and maturation on SLBs. In addition, our experiments and physical modeling show that cell adhesion on a fluid interface can be strongly modulated by the receptor-ligand affinity, in contrast with solid surfaces. These regulation levels might be used during selective adhesion of T- or B-cells, in which microtubules and dynein motors are known to be involved in clustering TCR and BCR receptors, respectively^78,79^.

## Data availability

Data supporting the findings of this study are available within the paper (and its supplementary information files). Numerical source data, raw microscopy images and Fiji/ImageJ macros for the detection of integrin clusters and membrane tubes are available upon reasonable request.

## Supporting information

Supplementary material Theory

Supplementary material video annotations

## Acknowledgments

We thank all laboratory members and collaborators, particularly Jay Groves (UC Berkeley) and Julien Pernier (Institute for Integrative Biology of the Cell) for their help in production of bilayers, Luke Lavis (Janelia Farm) for providing fluorophores, David Calderwood (Yale University) for providing MEF cell lines, Christof Hauck (University of Konstanz), Simon De Beco (Paris Diderot University) and Danijela Vignjevic (Institut Curie) for providing genetic constructs used in the study, Stephanie Miserey-Lenkei (Institut Curie) for the help in immunofluorescence experiments, Rémi Fert and Eric Nicolau for the help at the mechanical workshop, Jost Enninga (Institut Pasteur), Anna Akhmanova, Ana-Suncana Smith (Friedrich-Alexander-Universität), Jean-Baptiste Manneville (Laboratoire Matière et Systèmes Complexes, Paris-Cité University), Kristine Schauer (Institut Gustave Roussy), Pierre Sens, Gaelle Boncompain, Alexandre Baffet, Ryszard Wimmer (all Institut Curie) for insightful discussions. We also acknowledge the experimental support of the Molecular Biology and Cells platform at Institut Curie.

This work was supported by the Institut Curie, the Collège de France, the Institut National de la Santé et de la Recherche Médicale (Inserm) and the Centre National de la Recherche Scientifique (CNRS). We further acknowledge the Nikon Imaging Centre at Institut Curie-CNRS, member of the French National Research Infrastructure France-BioImaging (ANR10-INSB-04). O.M. was funded by the European Union’s Horizon 2020 research and innovation program under the Marie Skłodowska-Curie grant agreement No 666003, the ARC foundation, the Labex CelTisPhyBio (Grant ANR-11-LABX-0038, ANR-10-IDEX-0001-02). R.M.A. acknowledges funding from Fondation pour la Recherche Médicale (FRM Postdoctoral Fellowship). P.B. and J.F.J. are members of the CNRS consortium Approches Quantitatives du Vivant, the Labex CelTisPhyBio (ANR-11-LABX0038) and Paris Sciences et Lettres (ANR-10-IDEX-0001-02).

## Author contributions

O.M., P.B. and G.T.V.N. conceived the study. P.B. and G.T.V.N. acquired funding and equally contributed to this work. O.M. designed and performed experiments, analyzed data, and wrote the original manuscript. A.S.M. and O.M. developed image analysis programs for integrin cluster. F.D.F, J.M. and O.M. performed cloning, expression, purification and labeling of Invasin. A.B. and F.T generated lentiviral constructs and cell lines used in the study. J.F.J. supervised the theoretical work performed and described in the supplementary material by R.M.A. and M.T.C. who equally contributed to this work. O.M., G.T.V.N. and P.B. reviewed and edited the final manuscript. All authors approved the final manuscript prior to submission.

## Supplementary video V1

Brightfield time-lapse imaging of a trembling MEF cell on Invasin-SLB. Frames were acquired at an interval of 10 sec and video playback speed is at 7 fps. Time in min:sec. Scale bar: 5 µm.

## Supplementary video V2

Brightfield time-lapse imaging of an adherent MEF cell on Invasin-SLB. Frames were acquired at an interval of 10 sec and video playback speed is at 7 fps. Time in min:sec. Scale bar: 5 µm.

---

**Supplementary Figure S1.**
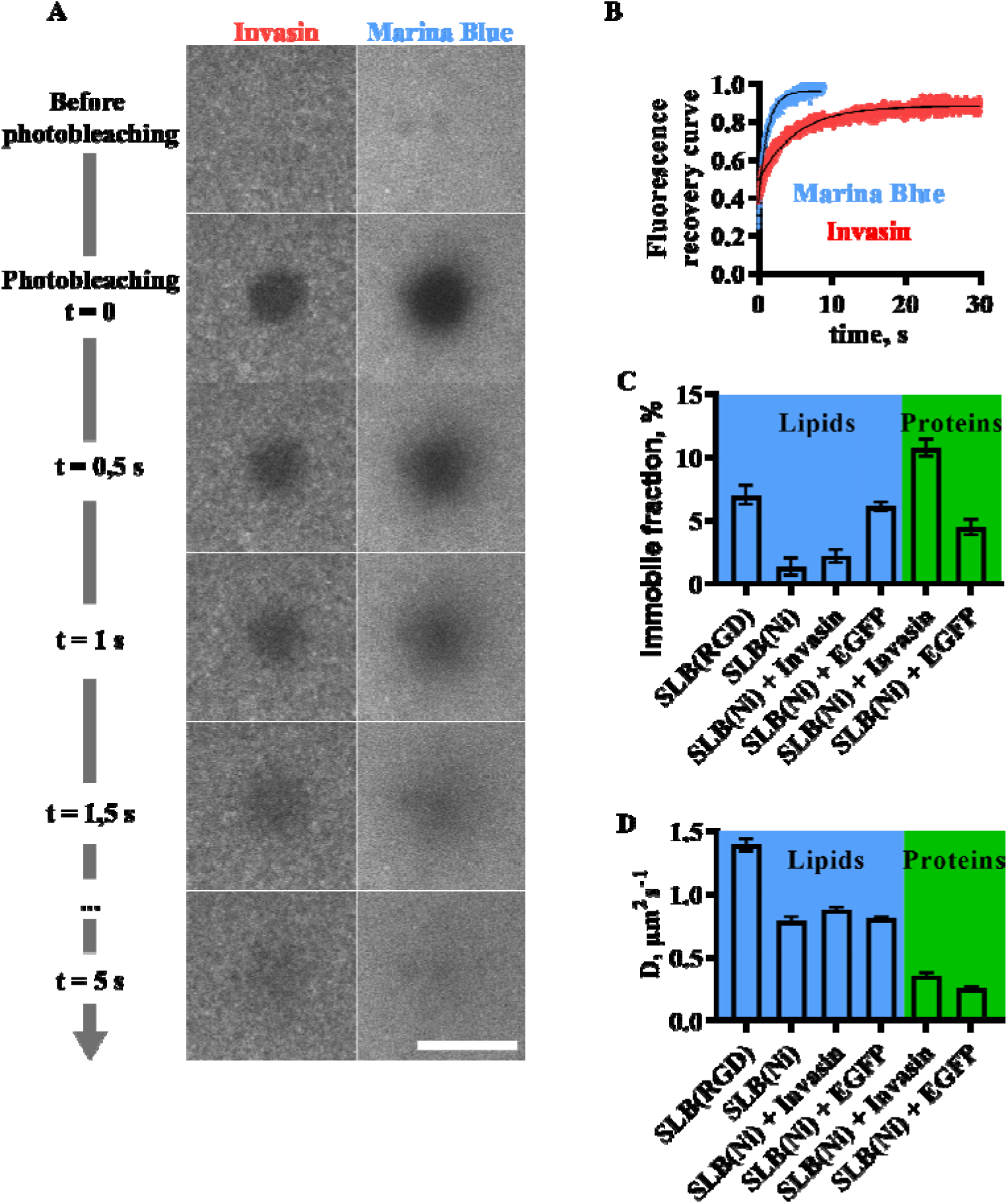
FRAP analysis of SLB fluidity. For each sample we have measured 10 FRAP curves in 2 independent experiments from a photobleached circular zone of approximately 20 µm^2^. A) Illustration of fluorescence recovery after photobleaching. Scale bar: 10 µm. B) Fluorescence recovery curves normalized by the fluorescence of the neighboring unbleached regions (blue – lipids labeled with Marina Blue, red – Invasin labeled with JF549). C-D) Different types of SLB surfaces: C) Immobile fractions of proteins and lipids. D) Diffusion coefficients of proteins and lipids.

**Supplementary Figure S2.**
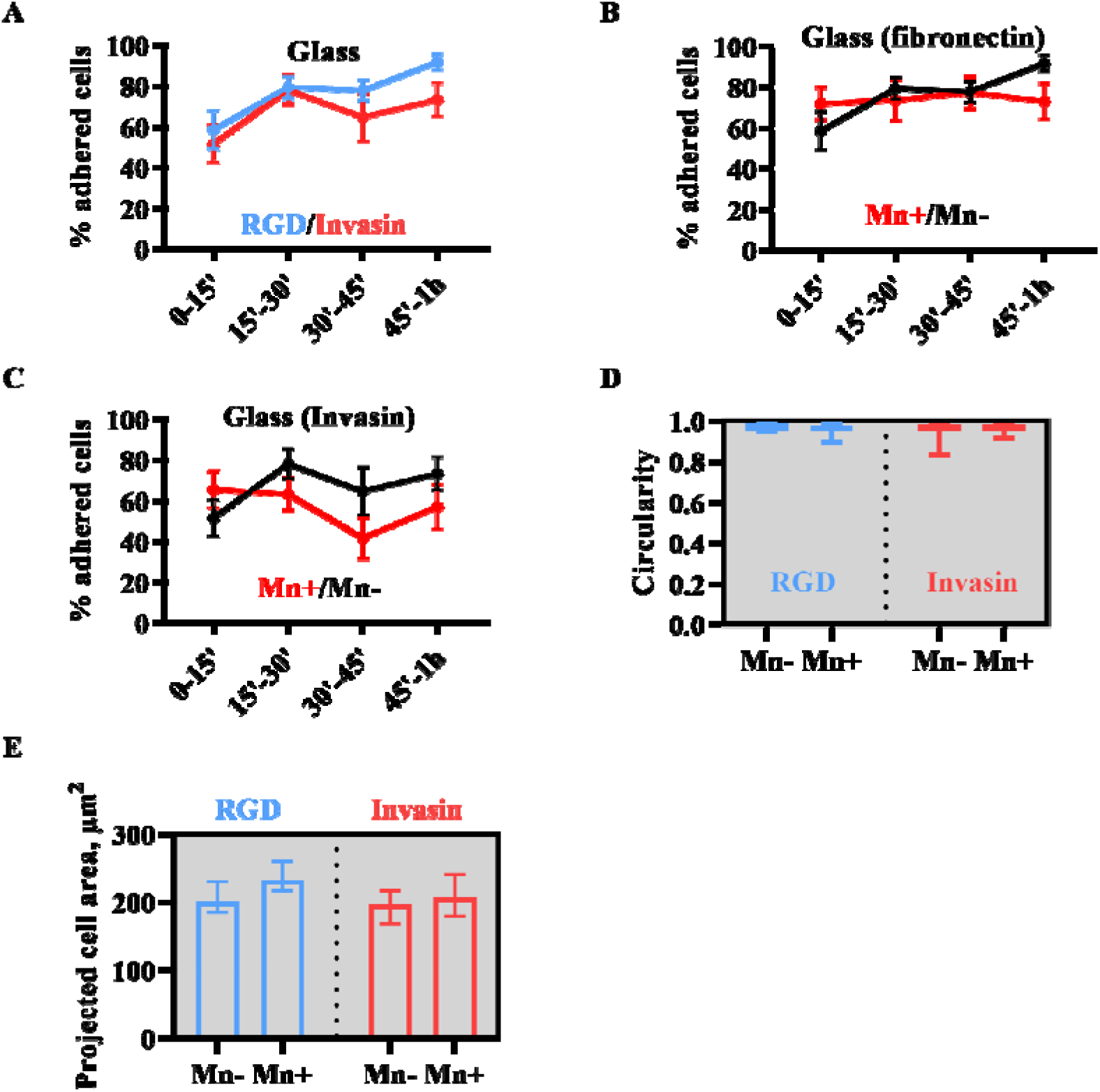
Percentage of “adherent” (“non-trembling”) MEF cells on glass over time and morphology of “trembling” MEF cells on SLBs. A-C) Time evolution of fractions of adherent cells on functionalized glass. For fibronectin (Mn-) each data point represents between 29 and 63 cells studied in at least 3 independent biological experiments (N=3). For fibronectin (Mn+): between 19 and 32 cells studied in at least N=3. For Invasin (Mn-): between 17 and 32 cells studied in at least N=3. For Invasin (Mn+): between 21 and 30 cells studied in at least N=3. Line scatter plots, mean, SEM. A) With fibronectin (RGD, blue) and Invasin (red) in the absence of Mn^2+^. B) With fibronectin (RGD) in the presence (Mn+, red) and in the absence (Mn-, black) of Mn^2+^. C) With Invasin in the presence (Mn+, red) and in the absence (Mn-, black) of Mn^2+^. D-E) Cell morphology of “trembling” MEF on SLBs during 1h after seeding to chambers. Data from 44 cells studied in at least 3 independent biological experiments (N=3) for RGD-SLB (Mn-); 35 cells studied in at least N=3 for RGD-SLB (Mn+); 57 cells studied in at least 3 independent biological experiments (N=3) for Invasin-SLB (Mn-); 46 cells studied in at least N=3 for Invasin-SLB (Mn+). D) Cell circularity. Box plots. E) Projected cell area. Bar plots, mean, SEM.

**Supplementary Figure S3.**
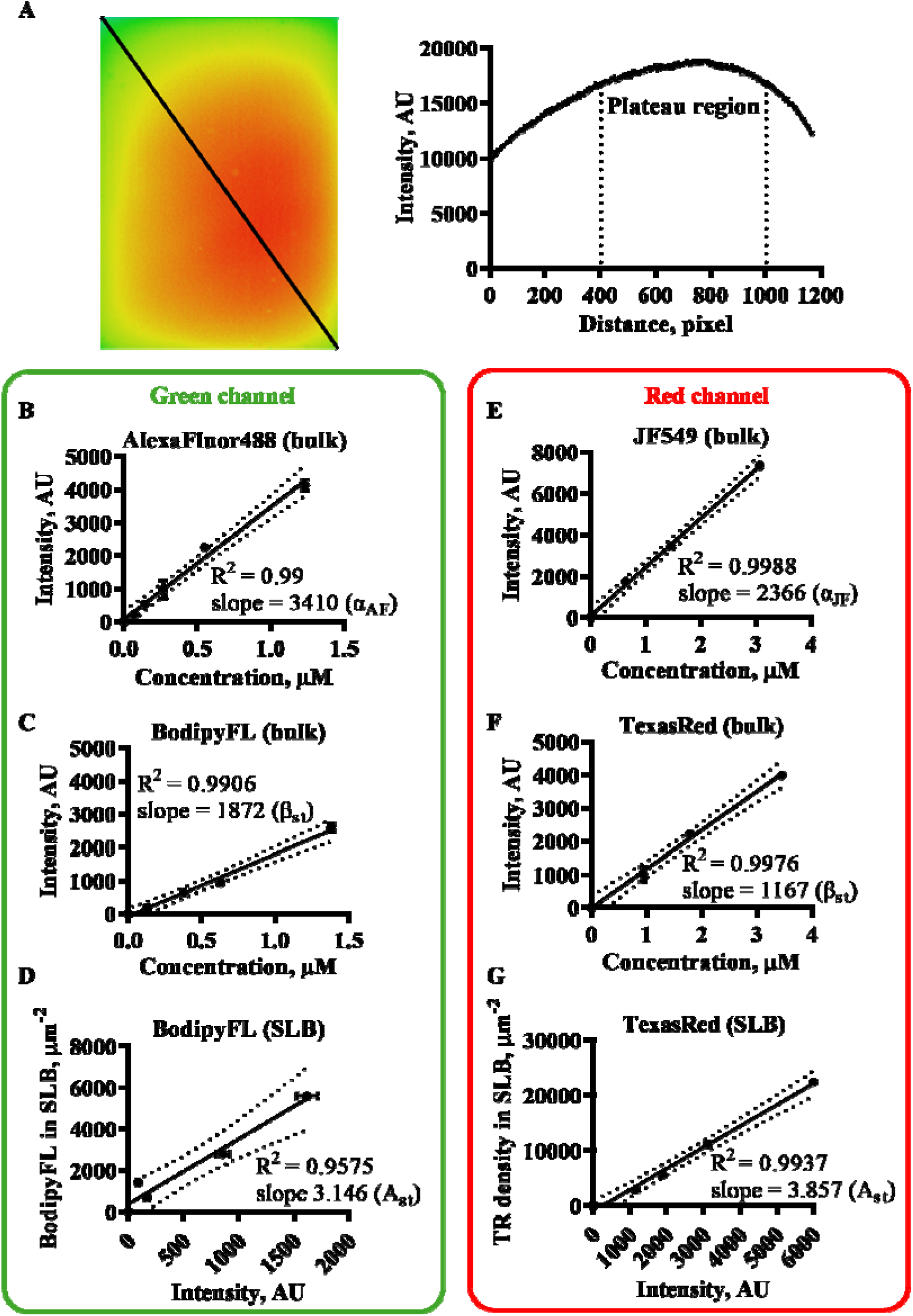
Protein density calibration using fluorescence in bulk and on SLBs (Methods). A) Illumination field (left) and profile along the black line (right) revealed by imaging AlexaFluor 488 dye in bulk solution. B-D) AlexaFluor488 fluorescence calibration: B) Fluorescence intensity – concentration curve for AlexaFluor488 in bulk. C) Fluorescence intensity – concentration curve for BodipyFL in bulk. D) Concentration – fluorescence intensity curve for BodipyFL on SLB. E-G) JF549 fluorescence calibration: E) Fluorescence intensity – concentration curve for JF549 in bulk. F) Fluorescence intensity – concentration curve for TexasRed in bulk. G) Concentration – fluorescence intensity curve for TexasRed on SLB.

**Supplementary Figure S4.**
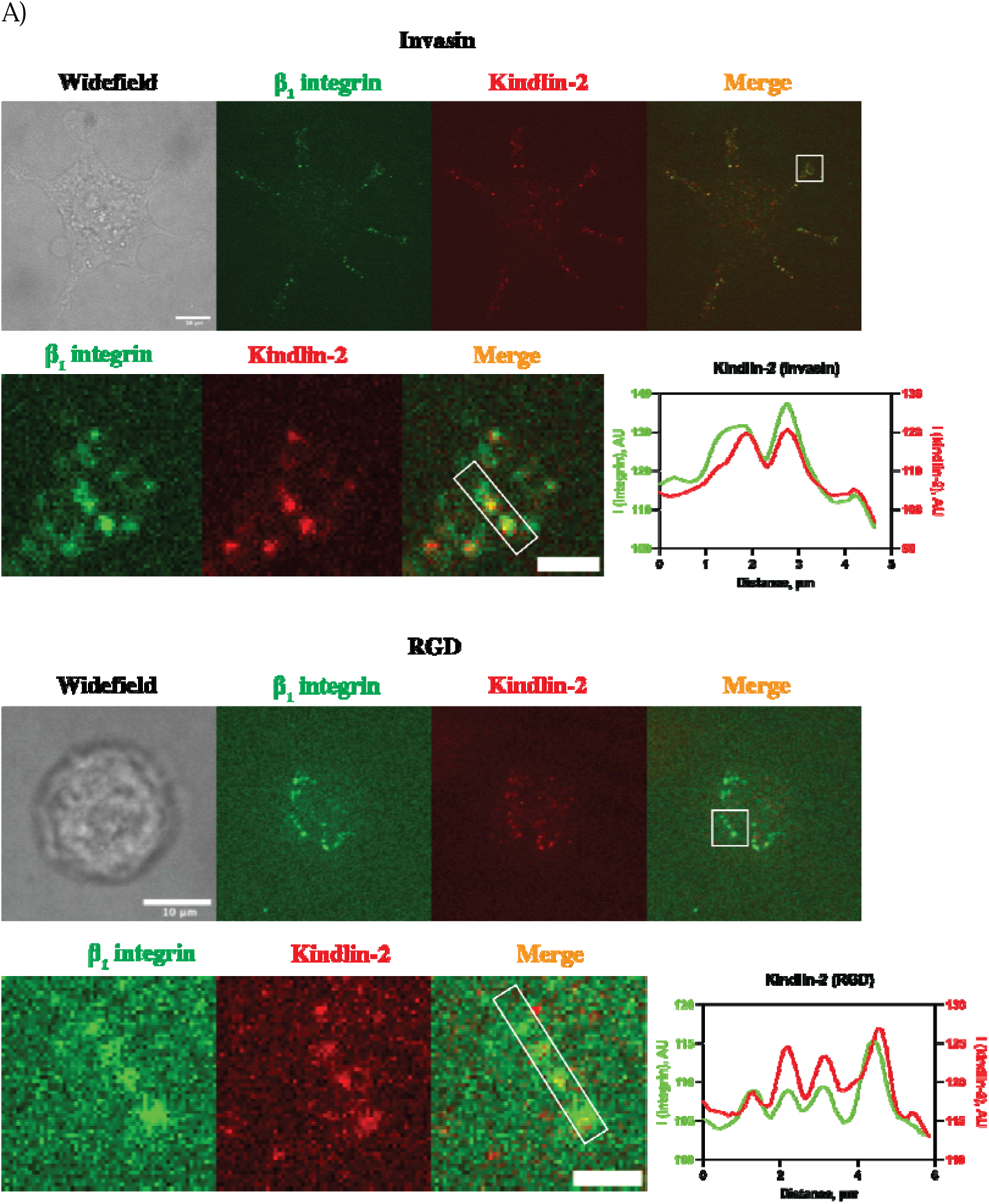
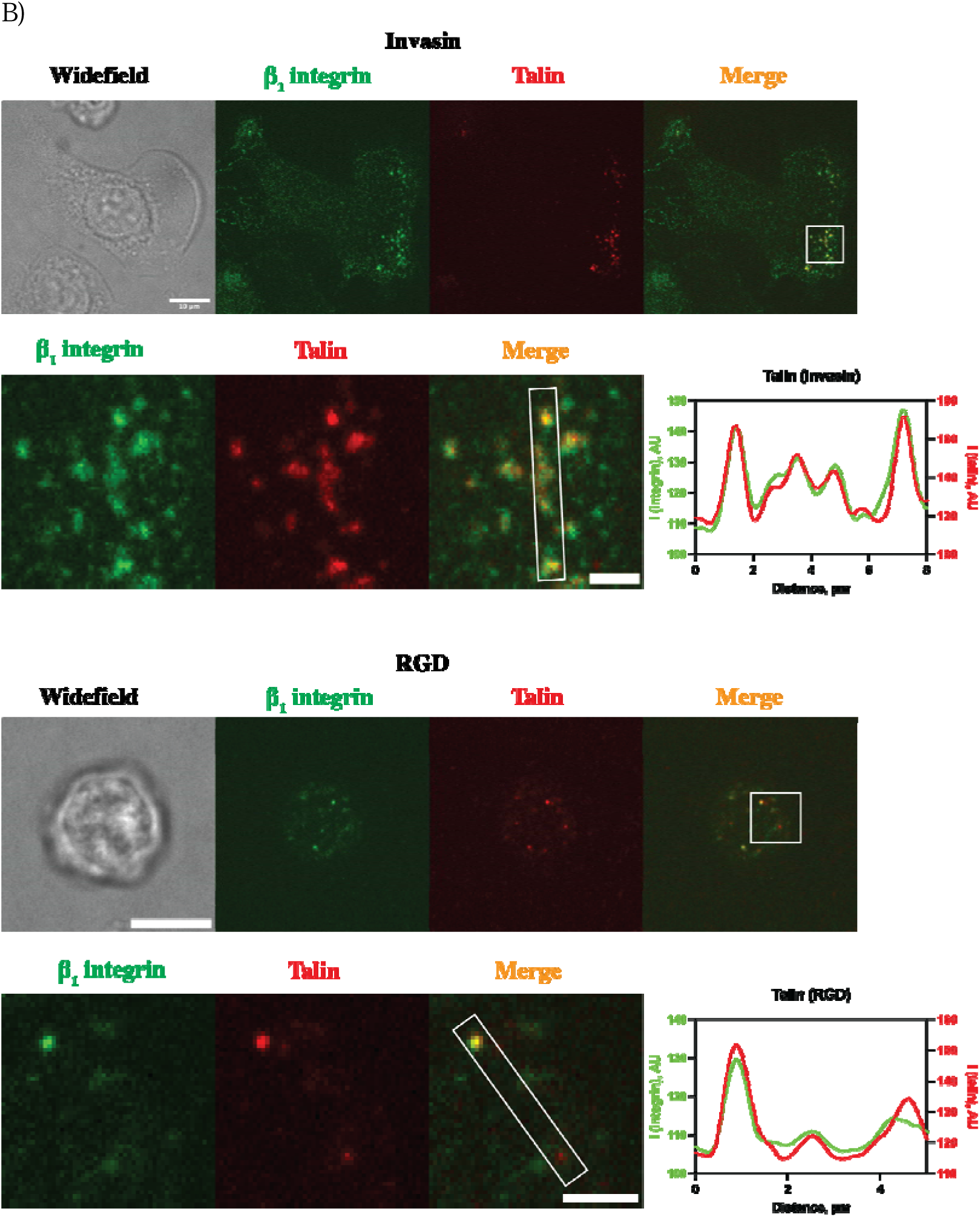
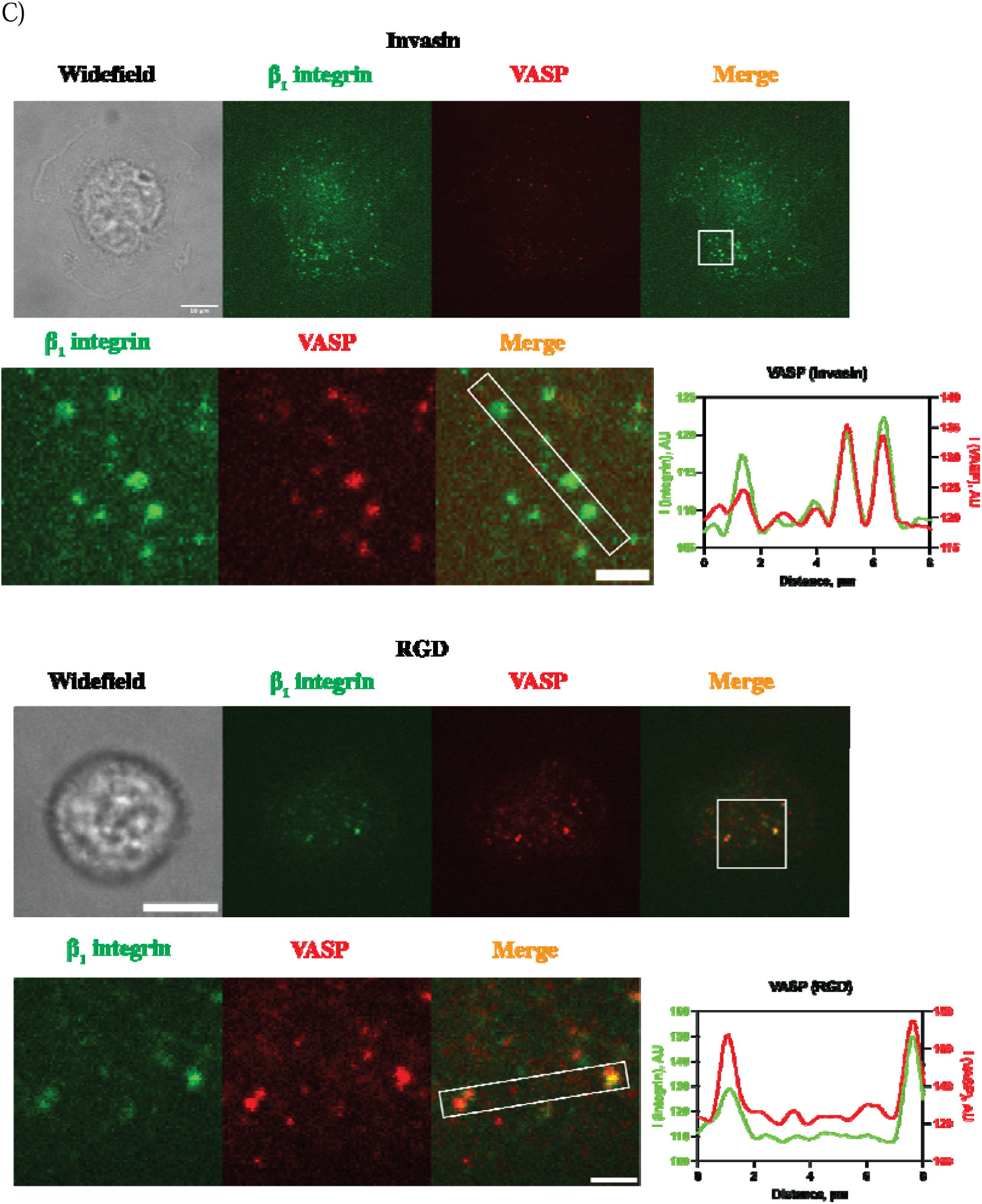
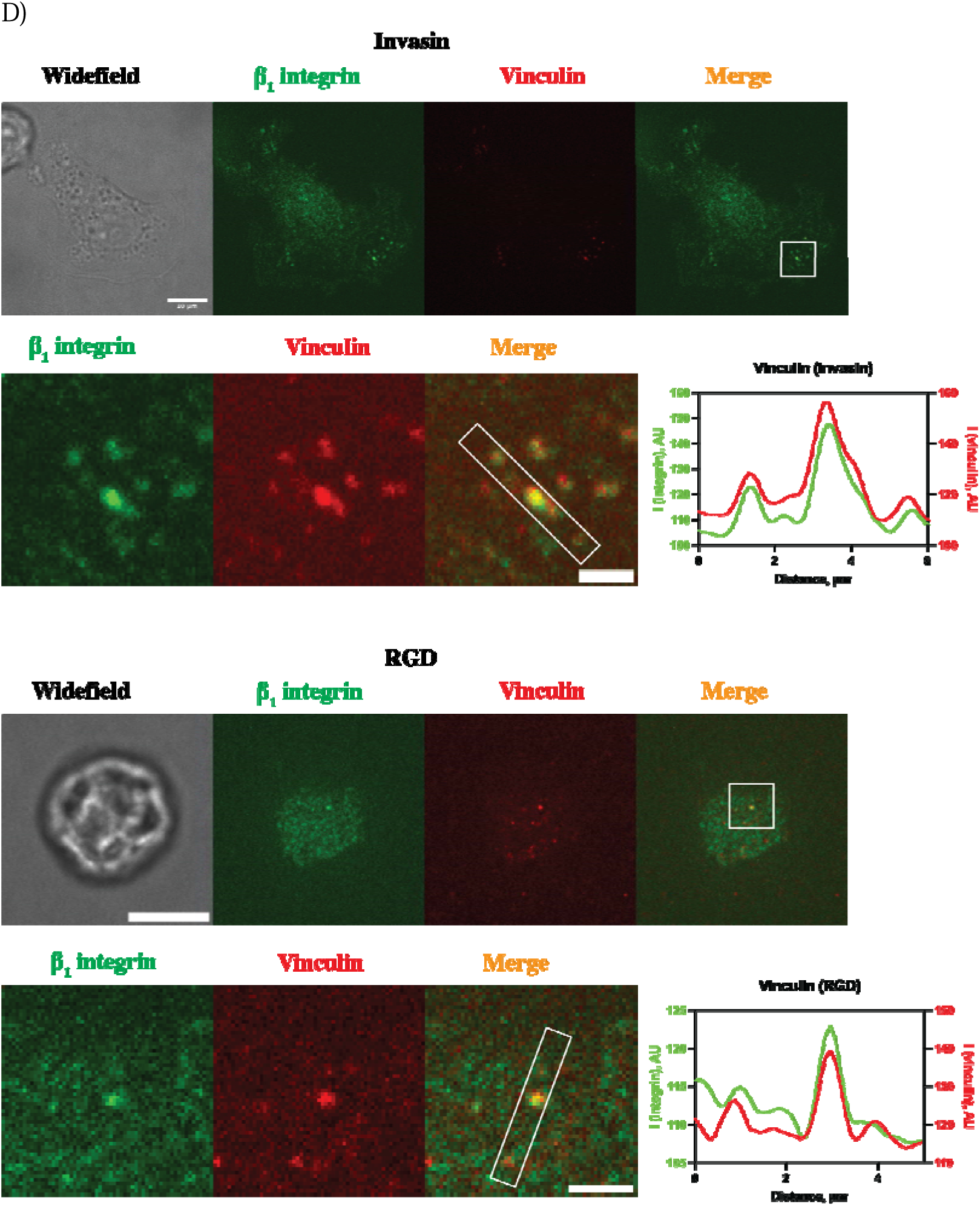
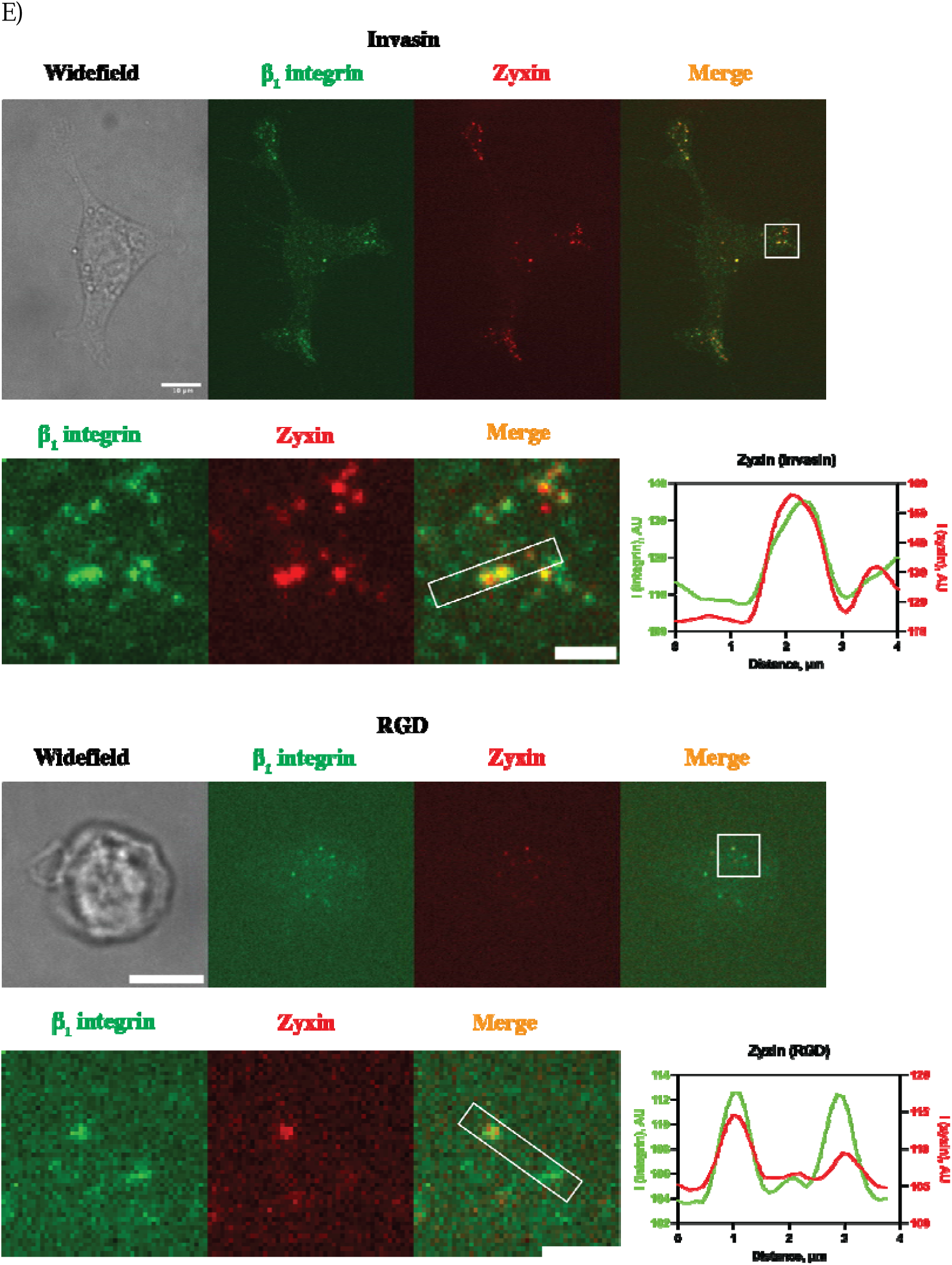
Series of figures corresponding to the data shown in Figure 3B illustrating the organization of integrin and other focal adhesion proteins (kindlin-2 (A), talin (B), VASP (C), vinculin (D), zyxin (E)) in clusters for cells adhering on SLBs in the presence of Mn^2+^. **Main panels**: Widefield image of the cell (grey, left) and fluorescence multi-channel images showing β_1_-integrin (green) and the respective focal adhesion protein (red) in cells adhering on SLBs coated with Invasin (top) and RGD (bottom). Images are taken at the focal plane of the SLB. **Bottom panels**: Left: zooms corresponding to the white squares in the main panels showing β_1_-integrin and focal adhesion proteins in the clusters. Right: β_1_-integrin and focal adhesion protein intensity profiles along the lines (white rectangles in the zoomed panels). Scale bars: 10 µm (main panels; all cells on Invasin; kindlin-2, talin, VASP, vinculin, zyxin cells on RGD); 2 µm (zoomed panels).

**Supplementary Figure S5.**
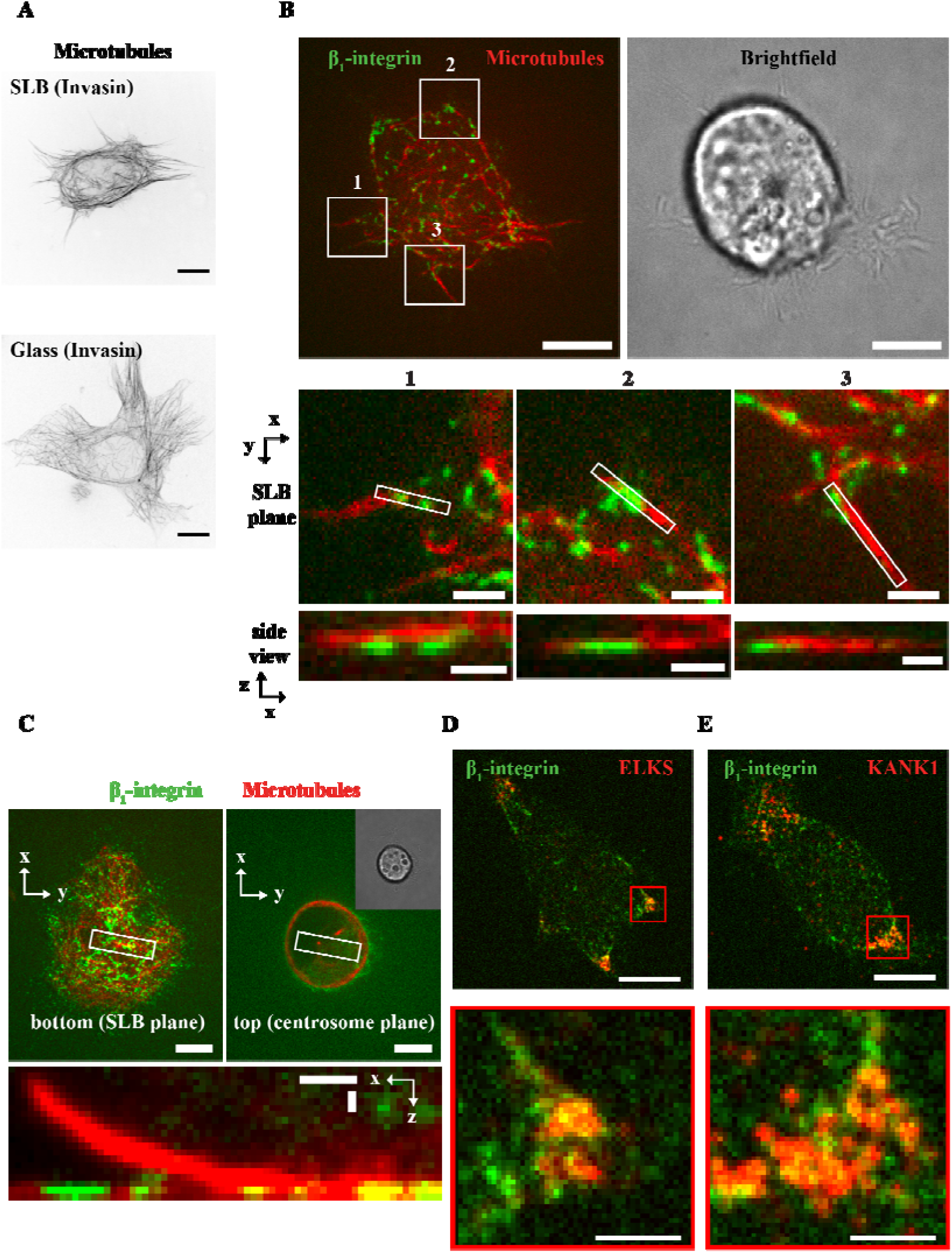
Microtubule organization, association with peripheral β_1_-integrin clusters and colocalization of microtubule adaptor proteins with β_1_-integrin clusters in MEF cells. A-C) Microtubules stained with SiR-tubulin. A) Distributions of microtubules in the adhesion plane of MEF cells on SLB (top) and glass (bottom) coated with Invasin. Scale bars: 10 µm. B) Main panels: distributions of β_1_-integrin clusters (green) and microtubules (red) at the SLB plane (top left) and brightfield image of a MEF cell on an Invasin-SLB. Scale bars: 10 µm. The 3 zoom panels correspond to the white frames in the main panel (left) and show the association of peripheral β_1_-integrin clusters with microtubules at the cell border in the xy and xz (orthogonal to the SLB) planes. Scale bars: 2 µm (zoomed panel xy); 1 µm (zoomed panel xz). C) Main panels: confocal images showing microtubules and β_1_-integrin clusters in the SLB plane (top left) and in the centrosome plane (top right) for a MEF cell on an Invasin-SLB. Scale bars: 10 µm. Bottom zoom panel (corresponding to the white frames in the main panels): orthogonal (xz) section to the SLB containing a centrosomal microtubule. Scale bars: 1 µm (zoomed panel xy); 1 µm (zoomed panel xz). D-E) ELKS (D) and KANK1 (E) (both red) colocalize with β_1_-integrin clusters (green) in MEF on Invasin-SLBs. Main panels (top) and zoomed panels (bottom) corresponding to the red frames in the main panels are shown. Scale bars: 10 µm (main panels); 2 µm (zoomed panels).

**Supplementary Figure S6.**
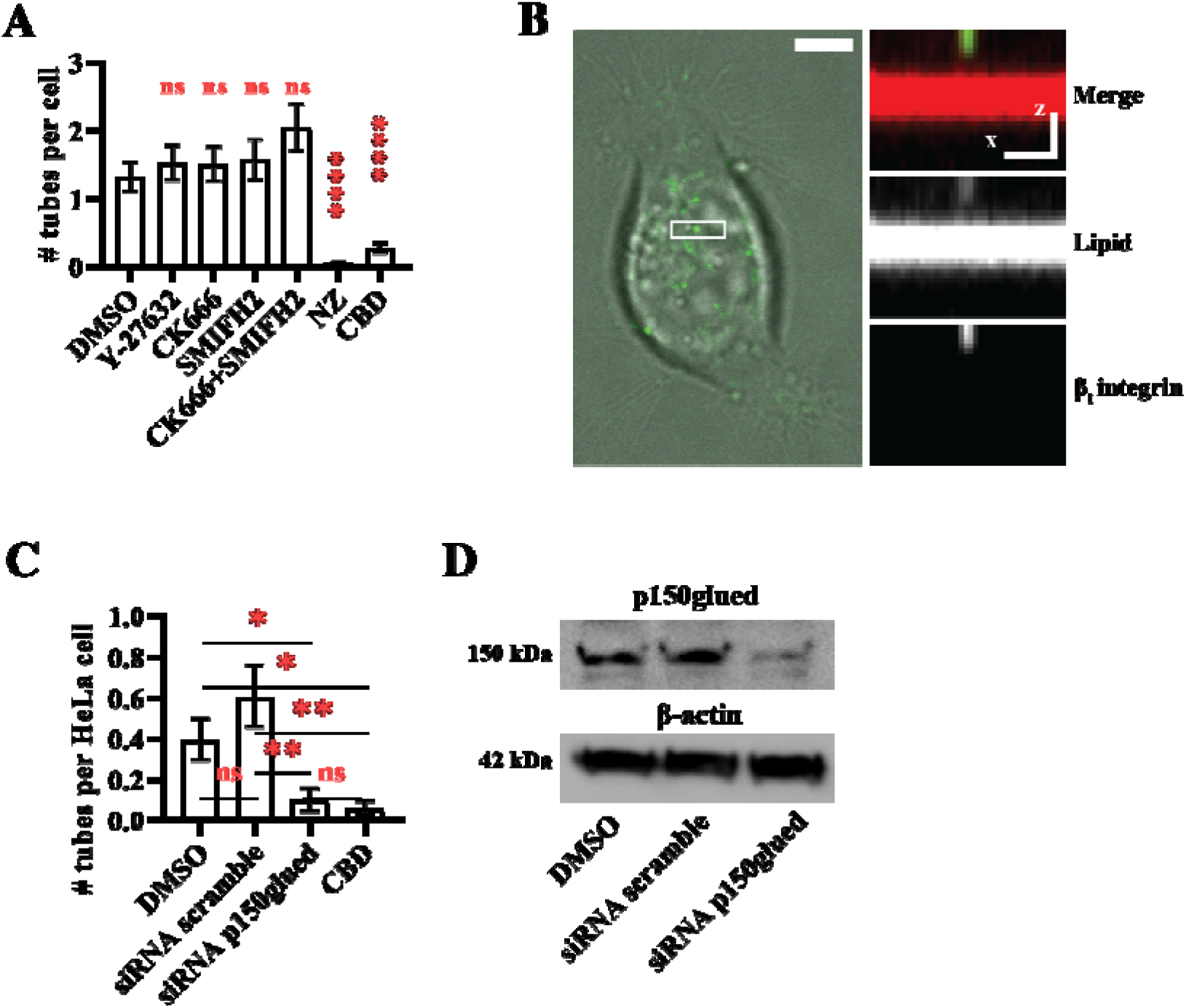
Number of tubes per cell (MEF and HeLa) under different treatments (drug inhibitors, siRNA) on Invasin-SLBs. A) Number of tubes in **MEF cells** treated with drug inhibitors and siRNA in the presence of Mn^2+^ between 0 – 1h after seeding on Invasin-SLBs. Bar plots: mean, SEM. Data from 78 cells, N=4 with DMSO; 104 cells, N=4 with CK666; 71 cells, N=3 with SMIFH2; 58 cells, N=3 with CK666+SMIFH2; 72 cells, N=3 with Y27632; 86 cells, N=3 with Nocodazole (NZ); 69 cells, N=4 with Ciliobrevin D (CBD). B) Image of an HeLa cell: brightfield (grey, left panel) and zoomed panel (right) showing a xz cross-section along the region corresponding to the white rectangle on the left. Bottom to top: β_1_-integrin with Halotag-Alexa488 (green), SLB and membrane tube labelled with a TR-DHPE lipid (red) and a merge of both channels. Scale bars: 5 µm (left), 1 µm (right, zoom). C) Number of tubes in **HeLa cells** treated with drug inhibitors and siRNA in the presence of Mn^2+^ between 0 – 1h after seeding on Invasin-SLBs. Bar plots: mean, SEM. Data from 58 cells, N=3 for DMSO; 56 cells, N=3 for siRNA scramble; 61 cells, N=4 for siRNA p150glued and 50 cells, N = 4 for CBD. D) Representative Western Blots showing a 77% reduction in the expression of p150glued subunit of dynactin in HeLa cells due to siRNA knockdown. Lysates of HeLa cells were prepared 48h after transfection with siRNA. p150glued (top) and β-actin (bottom) protein expression levels were analyzed by Western blotting using antibodies recognizing these proteins.

**Supplementary Figure S7.**
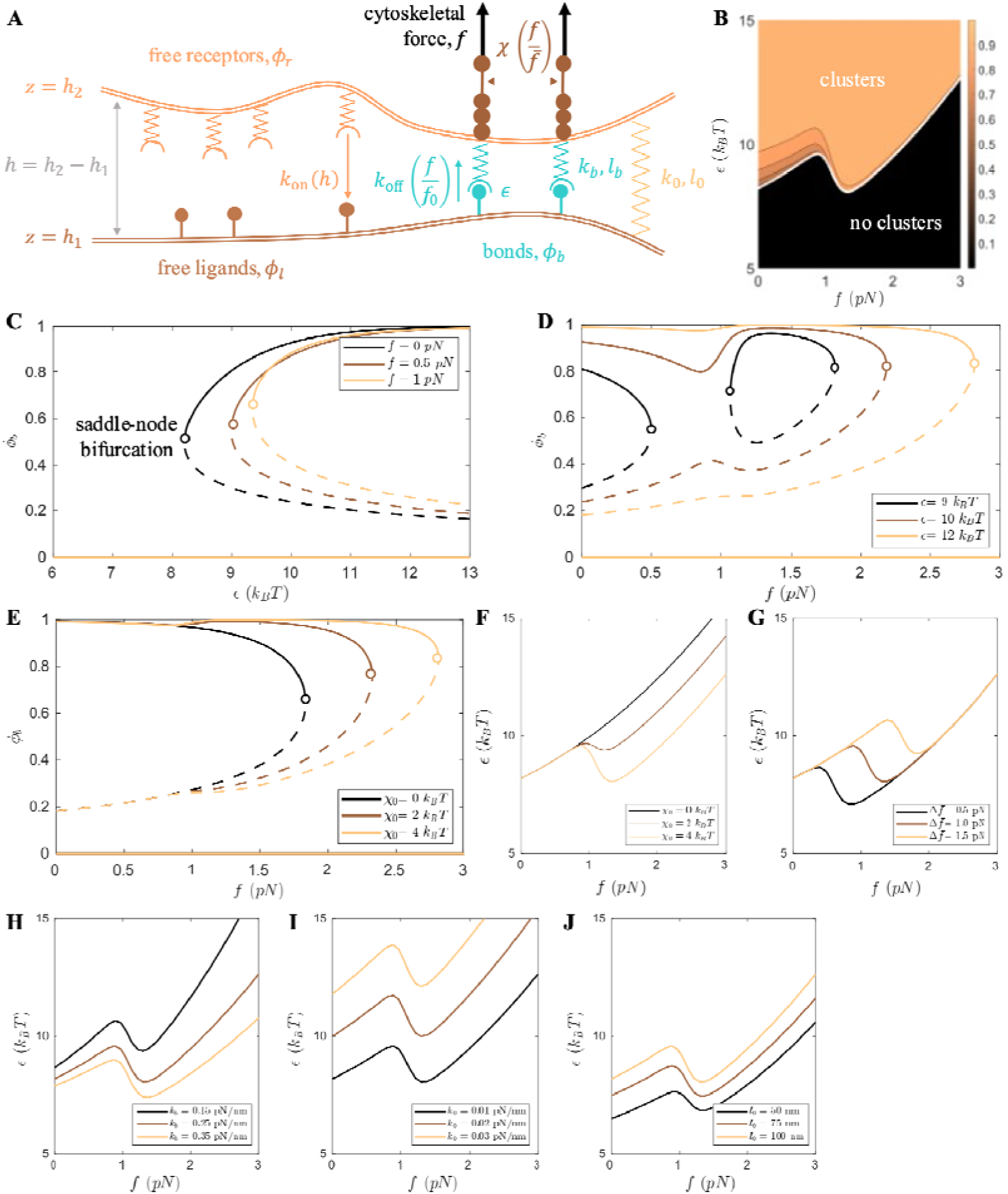
Theoretical predictions of integrin clustering as phase separation. A) Schematic illustration of the model. Free receptors and ligands form bound complexes (bonds) at a rate which depends on the distance between the cell membrane and the SLB,. The binding process is characterised by a gain in binding energy,. Once formed, a bond dissociates at a rate that increases when the bond is under force. The cytoskeleton applies a constant force, to each bond, which can stretch the adaptor proteins (force scale) or even break the bond (force scale). The interaction strength, describes the effective attractive interaction between bonds. The bonds are modelled as harmonic springs with rest length and spring constant, while the short-range interaction between cell membrane and SLB is also modelled as a spring of rest length and spring constant. B-C) Clustering as phase separation. B) Phase diagram of the system against the cytoskeletal force per bond, *f*, and the binding energy, *∈*. Above a critical binding energy (white line), the system can sustain two distinct bond concentrations at steady state, indicating that clusters can form. The color indicates the highest sustainable bond density. C) Bifurcation diagram of the steady state bond concentration against the binding energy, *∈*. The dilute phase (solid line, *Φ_b_*≈0) always exists, but a dense phase (solid line, *Φ_b_*→1) appears above a critical binding energy through a saddle-node bifurcation. The middle solution (dashed line) is always unstable. D-E) Clustering under force. The clusters can sustain larger forces (i.e. the dense phase persists until a larger cytoskeletal force per bond) for (D) higher affinity ligands (increasing *ε*) or (E) stronger attractive interaction through the adaptor proteins (increasing *χ*_0_). In (E), the value of the binding energy is *∈* = 12*k_B_T*. F-J) Dependence of phase diagram on model parameters. Each subfigure depicts the critical binding energy for clustering (white line in B) for several values of (F) interaction strength of activated bonds, *χ*_0_, (G) force required to activate the bonds, Δ*f̄*, (H) spring constant of the receptors/bonds, *k_b_*, (I) spring constant of the glycocalyx, *k*_0_, and (J) size of the glycocalyx, *l*_0_.

**Supplementary Figure S8.**
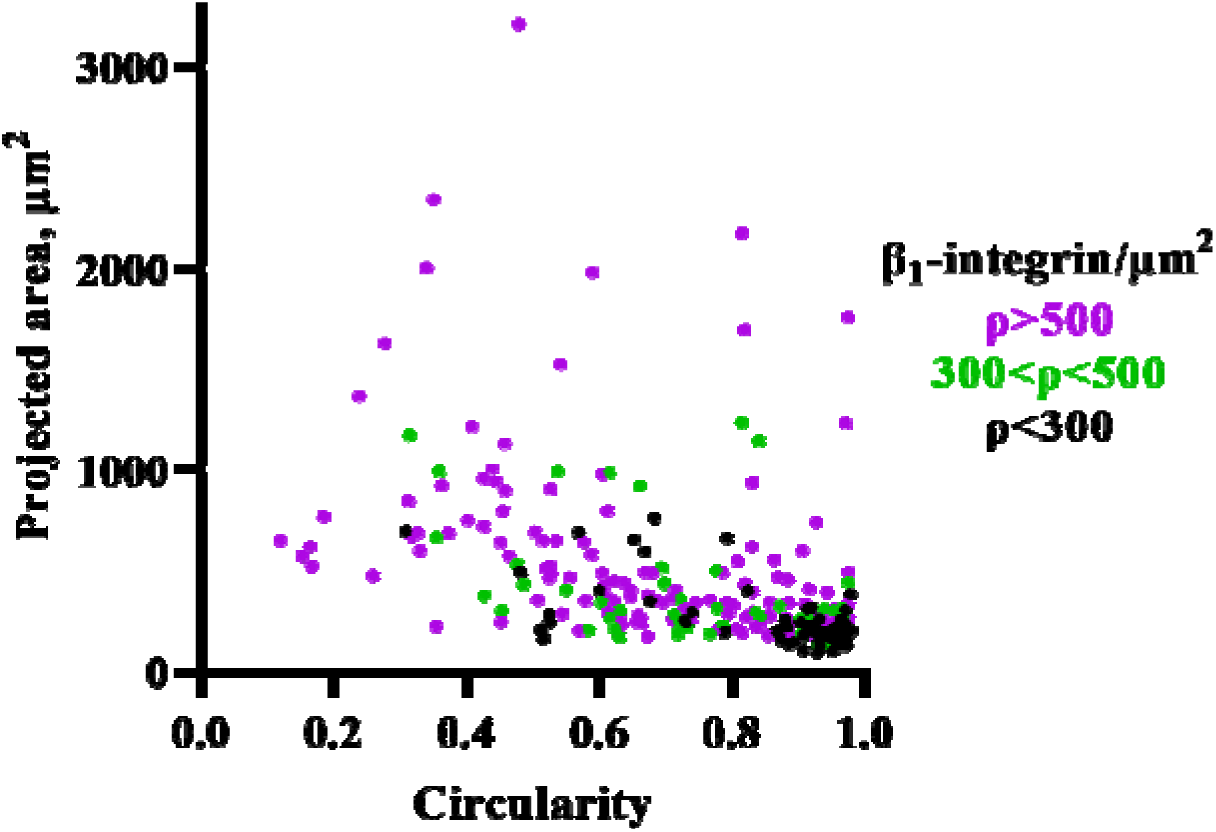
Correlation between the maximal β_1_-integrin density in the clusters (ρ) and the projected area and the circularity of Mn^2+^-treated MEF cells on Invasin-SLBs between 0 – 1h after seeding to chambers. Cloud plot of 3 subpopulations of cells with different integrin densities (ρ > 500, 300 < ρ < 500 and ρ < 300 integrins/μm^2^). It shows the correlation between cell morphologies (projected area and circularity) and protein density. Data from 157 cells studied in at least 3 independent biological experiments (N=3) for ρ > 500; 61 cells studied in at least N=3 for 300 < ρ < 500 and 87 cells studied in at least N=3 for ρ < 300 integrins/μm^2^.

**Supplementary Figure S9.**
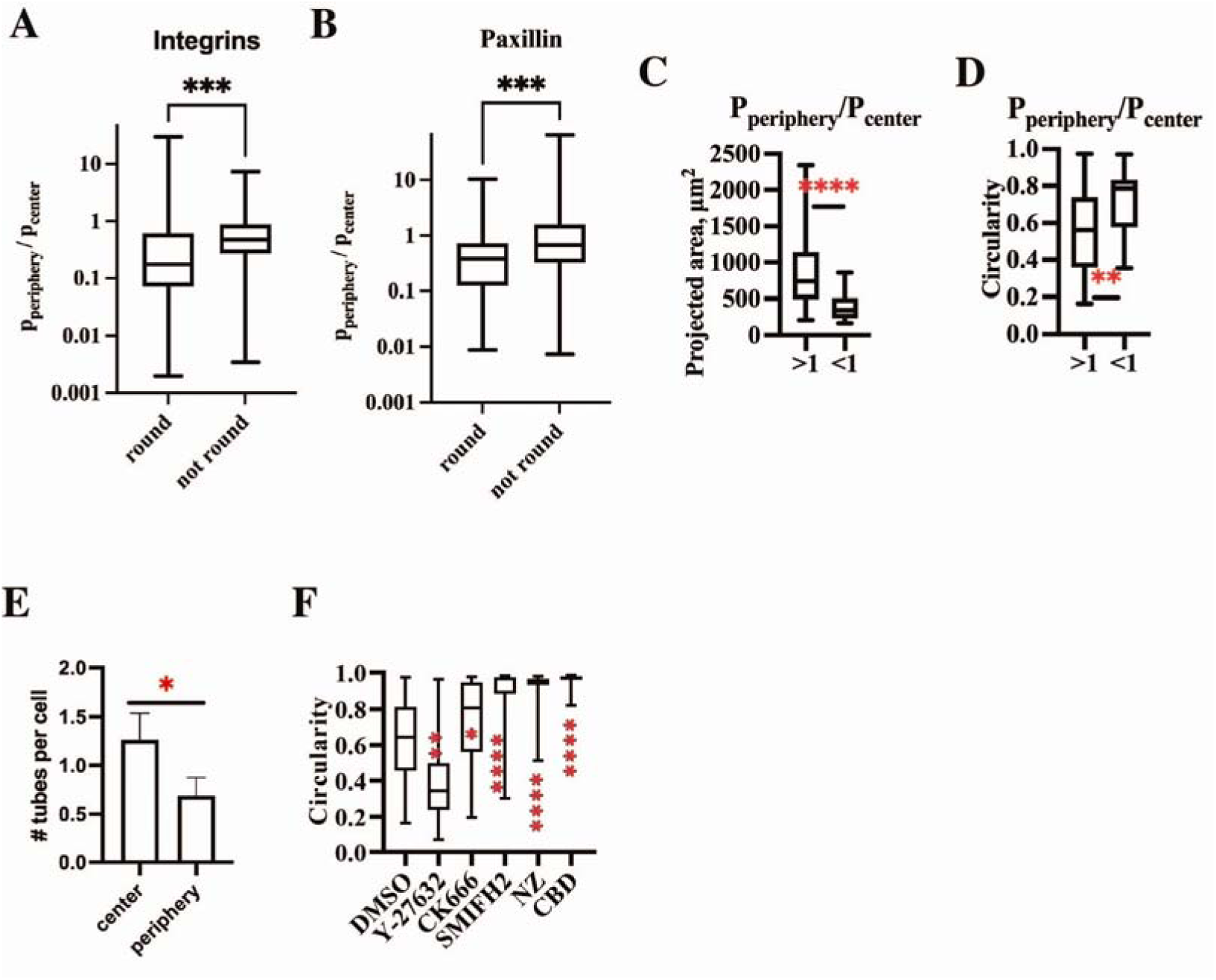
Peripheral vs. central localization of adhesion clusters in cells on Invasin-SLBs. A) Box plots of the ratios of area fractions of β_1_-integrin clusters in the periphery zone (P_periphery_) to the center zone (P_center_) for round (circularity > 0.8) and non-round (circularity < 0.8). Cells were allowed to adhere on Invasin-coated SLBs for 45-60 min in the presence of Mn^2+^. N=3, 78 cells. Mann-Whitney statistical test. B) Box plots of the ratios of area fractions of paxillin clusters in the periphery zone (P_periphery_) to the center zone (P_center_) for round (circularity > 0.8) and non-round (circularity < 0.8). Cells were allowed to adhere on Invasin-coated SLBs for 45-60 min in the presence of Mn^2+^. N=3, 119 cells. Mann-Whitney statistical test. C) Projected area of cells with higher β_1_-integrin cluster areas at the cell periphery (P_periphery_/ P_center_>1) or at the cell body (P_periphery_/ P_center_<1). Box plot. D) Cell circularity of cells with higher β_1_-integrin cluster areas at the cell periphery (P_periphery_/ P_center_>1) or at the cell body (P_periphery_/ P_center_<1). Box plot. E) Number of detected tubes at the periphery and in the center in spread MEF cells (projected area > 450 µm^2^) adhering on Invasin-SLBs. Bar plot, mean, SEM. N=3, 42 cells. Mann-Whitney statistical test. F) Circularity of cells treated with drugs. Box plot. Data from 78 cells, N=4 with DMSO; 104 cells, N=4 with CK666; 71 cells, N=3 with SMIFH2; 58 cells, N=3 with CK666+SMIFH2; 72 cells, N=3 with Y27632; 86 cells, N=3 with Nocodazole (NZ); 69 cells, N=4 with Ciliobrevin D (CBD).

## Bibliography

1. Huttenlocher, A. & Horwitz, A. R. Integrins in Cell Migration. Cold Spring Harb Perspect Biol 3, a005074 (2011).

2. Vogel, V. & Sheetz, M. P. Cell fate regulation by coupling mechanical cycles to biochemical signaling pathways. Curr Opin Cell Biol 21, 38–46 (2009).

3. Bökel, C. & Brown, N. H. Integrins in Development: Moving on, Responding to, and Sticking to the Extracellular Matrix. Developmental Cell 3, 311–321 (2002).

4. Discher, D. E., Janmey, P. & Wang, Y.-L. Tissue cells feel and respond to the stiffness of their substrate. Science 310, 1139–1143 (2005).

5. Iskratsch, T., Wolfenson, H. & Sheetz, M. P. Appreciating force and shape — the rise of mechanotransduction in cell biology. Nat Rev Mol Cell Biol 15, 825–833 (2014).

6. Winograd-Katz, S. E., Fässler, R., Geiger, B. & Legate, K. R. The integrin adhesome: from genes and proteins to human disease. Nat Rev Mol Cell Biol 15, 273–288 (2014).

7. Sun, Q., Hou, Y., Chu, Z. & Wei, Q. Soft overcomes the hard: Flexible materials adapt to cell adhesion to promote cell mechanotransduction. Bioactive Materials 10, 397–404 (2022).

8. Sun, Z., Guo, S. S. & Fässler, R. Integrin-mediated mechanotransduction. J Cell Biol 215, 445–456 (2016).

9. Choi, C. K. et al. Actin and alpha-actinin orchestrate the assembly and maturation of nascent adhesions in a myosin II motor-independent manner. Nat Cell Biol 10, 1039–1050 (2008).

10. Bershadsky, A. D. et al. Assembly and mechanosensory function of focal adhesions: experiments and models. Eur J Cell Biol 85, 165–173 (2006).

11. Yamada, K. M., Doyle, A. D. & Lu, J. Cell–3D matrix interactions: recent advances and opportunities. Trends in Cell Biology (2022) doi:10.1016/j.tcb.2022.03.002.

12. Huveneers, S. & de Rooij, J. Mechanosensitive systems at the cadherin-F-actin interface. J Cell Sci 126, 403–413 (2013).

13. Harjunpää, H., Llort Asens, M., Guenther, C. & Fagerholm, S. C. Cell Adhesion Molecules and Their Roles and Regulation in the Immune and Tumor Microenvironment. Frontiers in Immunology 10, (2019).

14. Rose, D. M., Alon, R. & Ginsberg, M. H. Integrin modulation and signaling in leukocyte adhesion and migration. Immunological Reviews 218, 126–134 (2007).

15. Galush, W. J., Nye, J. A. & Groves, J. T. Quantitative fluorescence microscopy using supported lipid bilayer standards. Biophys J 95, 2512–2519 (2008).

16. Yu, C., Law, J. B. K., Suryana, M., Low, H. Y. & Sheetz, M. P. Early integrin binding to Arg-Gly-Asp peptide activates actin polymerization and contractile movement that stimulates outward translocation. Proc Natl Acad Sci U S A 108, 20585–20590 (2011).

17. Yu, C. et al. Integrin-matrix clusters form podosome-like adhesions in the absence of traction forces. Cell Rep 5, 1456–1468 (2013).

18. Glazier, R. & Salaita, K. Supported lipid bilayer platforms to probe cell mechanobiology. Biochimica et Biophysica Acta (BBA) - Biomembranes 1859, 1465–1482 (2017).

19. Springer, T. A. Adhesion receptors of the immune system. Nature 346, 425–434 (1990).

20. Langereis, J. D. Neutrophil integrin affinity regulation in adhesion, migration, and bacterial clearance. Cell Adh Migr 7, 476–481 (2013).

21. Isberg, R. R. & Leong, J. M. Multiple β1 chain integrins are receptors for invasin, a protein that promotes bacterial penetration into mammalian cells. Cell 60, 861– 871 (1990).

22. Huet-Calderwood, C. et al. Novel ecto-tagged integrins reveal their trafficking in live cells. Nat Commun 8, 570 (2017).

23. Shattil, S. J., Kim, C. & Ginsberg, M. H. The final steps of integrin activation: the end game. Nat Rev Mol Cell Biol 11, 288–300 (2010).

24. Ni, H., Li, A., Simonsen, N. & Wilkins, J. A. Integrin activation by dithiothreitol or Mn2+ induces a ligand-occupied conformation and exposure of a novel NH2-terminal regulatory site on the beta1 integrin chain. J Biol Chem 273, 7981–7987 (1998).

25. Galbraith, C. G., Yamada, K. M. & Galbraith, J. A. Polymerizing actin fibers position integrins primed to probe for adhesion sites. Science 315, 992–995 (2007).

26. Price, L. S., Leng, J., Schwartz, M. A. & Bokoch, G. M. Activation of Rac and Cdc42 by integrins mediates cell spreading. Mol Biol Cell 9, 1863–1871 (1998).

27. Cavalcanti-Adam, E. A. et al. Lateral spacing of integrin ligands influences cell spreading and focal adhesion assembly. Eur J Cell Biol 85, 219–224 (2006).

28. Wiseman, P. W. et al. Spatial mapping of integrin interactions and dynamics during cell migration by image correlation microscopy. J Cell Sci 117, 5521–5534 (2004).

29. Kanchanawong, P. et al. Nanoscale architecture of integrin-based cell adhesions. Nature 468, 580–584 (2010).

30. Stehbens, S. & Wittmann, T. Targeting and transport: How microtubules control focal adhesion dynamics. Journal of Cell Biology 198, 481–489 (2012).

31. LaFlamme, S. E., Mathew-Steiner, S., Singh, N., Colello-Borges, D. & Nieves, B. Integrin and microtubule crosstalk in the regulation of cellular processes. Cell. Mol. Life Sci. 75, 4177–4185 (2018).

32. Bouchet, B. P. et al. Talin-KANK1 interaction controls the recruitment of cortical microtubule stabilizing complexes to focal adhesions. eLife 5, e18124 (2016).

33. Lansbergen, G. et al. CLASPs attach microtubule plus ends to the cell cortex through a complex with LL5beta. Dev Cell 11, 21–32 (2006).

34. Bennett, M. et al. Molecular clutch drives cell response to surface viscosity. Proceedings of the National Academy of Sciences 115, 1192–1197 (2018).

35. Glazier, R. et al. DNA mechanotechnology reveals that integrin receptors apply pN forces in podosomes on fluid substrates. Nat Commun 10, 4507 (2019).

36. Ghassemi, S. et al. Cells test substrate rigidity by local contractions on submicrometer pillars. Proceedings of the National Academy of Sciences 109, 5328–5333 (2012).

37. Rafiq, N. B. M. et al. A mechano-signalling network linking microtubules, myosin IIA filaments and integrin-based adhesions. Nat. Mater. 18, 638–649 (2019).

38. Seetharaman, S. et al. Microtubules tune mechanosensitive cell responses. Nat. Mater. 21, 366–377 (2022).

39. Bershadsky, A., Chausovsky, A., Becker, E., Lyubimova, A. & Geiger, B. Involvement of microtubules in the control of adhesion-dependent signal transduction. Curr Biol 6, 1279–1289 (1996).

40. Chang, Y.-C., Nalbant, P., Birkenfeld, J., Chang, Z.-F. & Bokoch, G. M. GEF-H1 Couples Nocodazole-induced Microtubule Disassembly to Cell Contractility via RhoA. MBoC 19, 2147–2153 (2008).

41. Komura, S. & Andelman, D. Adhesion-induced lateral phase separation in membranes. Eur. Phys. J. E 3, 259–271 (2000).

42. Bihr, T., Seifert, U. & Smith, A.-S. Nucleation of Ligand-Receptor Domains in Membrane Adhesion. Phys. Rev. Lett. 109, 258101 (2012).

43. Fenz, S. F. et al. Membrane fluctuations mediate lateral interaction between cadherin bonds. Nature Phys 13, 906–913 (2017).

44. Geiger, B. & Bershadsky, A. Exploring the neighborhood: adhesion-coupled cell mechanosensors. Cell 110, 139–142 (2002).

45. Schwarz, U. S. & Safran, S. A. Physics of adherent cells. Rev. Mod. Phys. 85, 1327–1381 (2013).

46. Braeutigam, A., Simsek, A. N., Gompper, G. & Sabass, B. Generic self-stabilization mechanism for biomolecular adhesions under load. Nat Commun 13, 2197 (2022).

47. del Rio, A. et al. Stretching single talin rod molecules activates vinculin binding. Science 323, 638–641 (2009).

48. Yao, M. et al. The mechanical response of talin. Nat Commun 7, 11966 (2016).

49. Brockman, J. M. et al. Mapping the 3D orientation of piconewton integrin traction forces. Nat Methods 15, 115–118 (2018).

50. Shakiba, D. et al. The Balance between Actomyosin Contractility and Microtubule Polymerization Regulates Hierarchical Protrusions That Govern Efficient Fibroblast-Collagen Interactions. ACS Nano 14, 7868–7879 (2020).

51. Dehmelt, L., Nalbant, P., Steffen, W. & Halpain, S. A microtubule-based, dynein-dependent force induces local cell protrusions: Implications for neurite initiation. Brain Cell Bio 35, 39–56 (2006).

52. Chen, B.-H., Tzen, J. T. C., Bresnick, A. R. & Chen, H.-C. Roles of Rho-associated Kinase and Myosin Light Chain Kinase in Morphological and Migratory Defects of Focal Adhesion Kinase-null Cells*. Journal of Biological Chemistry 277, 33857–33863 (2002).

53. Iskratsch, T. et al. FHOD1 is needed for directed Forces and Adhesion Maturation during Cell Spreading and Migration. Dev Cell 27, 545–559 (2013).

54. Burridge, K. & Wittchen, E. S. The tension mounts: Stress fibers as force-generating mechanotransducers. J Cell Biol 200, 9–19 (2013).

55. Marlin, S. D. & Springer, T. A. Purified intercellular adhesion molecule-1 (ICAM-1) is a ligand for lymphocyte function-associated antigen 1 (LFA-1). Cell 51, 813–819 (1987).

56. Elices, M. J. et al. VCAM-1 on activated endothelium interacts with the leukocyte integrin VLA-4 at a site distinct from the VLA-4/fibronectin binding site. Cell 60, 577–584 (1990).

57. Livne, A. & Geiger, B. The inner workings of stress fibers[−[from contractile machinery to focal adhesions and back. Journal of Cell Science 129, 1293–1304 (2016).

58. Tojkander, S., Gateva, G. & Lappalainen, P. Actin stress fibers – assembly, dynamics and biological roles. Journal of Cell Science 125, 1855–1864 (2012).

59. Bouchet, B. P. & Akhmanova, A. Microtubules in 3D cell motility. Journal of Cell Science 130, 39–50 (2017).

60. Rhee, S., Jiang, H., Ho, C.-H. & Grinnell, F. Microtubule function in fibroblast spreading is modulated according to the tension state of cell–matrix interactions. Proc Natl Acad Sci U S A 104, 5425–5430 (2007).

61. Stehbens, S. J. et al. CLASPs link focal adhesion-associated microtubule capture to localized exocytosis and adhesion site turnover. Nat Cell Biol 16, 561–573 (2014).

62. Hendricks, A. G. et al. Dynein Tethers and Stabilizes Dynamic Microtubule Plus-Ends. Curr Biol 22, 632–637 (2012).

63. Rosse, C. et al. Binding of dynein intermediate chain 2 to paxillin controls focal adhesion dynamics and migration. Journal of Cell Science 125, 3733–3738 (2012).

64. Redwine, W. B. et al. The human cytoplasmic dynein interactome reveals novel activators of motility. eLife 6, e28257 (2017).

65. Leduc, C., Campàs, O., Joanny, J.-F., Prost, J. & Bassereau, P. Mechanism of membrane nanotube formation by molecular motors. Biochimica et Biophysica Acta (BBA) - Biomembranes 1798, 1418–1426 (2010).

66. Gumí-Audenis, B. et al. Pulling lipid tubes from supported bilayers unveils the underlying substrate contribution to the membrane mechanics. Nanoscale 10, 14763–14770 (2018).

67. Gennerich, A., Carter, A. P., Reck-Peterson, S. L. & Vale, R. D. Force-induced bidirectional stepping of cytoplasmic dynein. Cell 131, 952–965 (2007).

68. Moreno-Layseca, P. et al. Cargo-specific recruitment in clathrin- and dynamin-independent endocytosis. Nat Cell Biol 23, 1073–1084 (2021).

69. Kalappurakkal, J. M. et al. Integrin Mechano-chemical Signaling Generates Plasma Membrane Nanodomains that Promote Cell Spreading. Cell 177, 1738–1756.e23 (2019).

70. Mayor, S., Parton, R. G. & Donaldson, J. G. Clathrin-Independent Pathways of Endocytosis. Cold Spring Harb Perspect Biol 6, a016758 (2014).

71. Cavalcanti-Adam, E. A. et al. Cell Spreading and Focal Adhesion Dynamics Are Regulated by Spacing of Integrin Ligands. Biophysical Journal 92, 2964–2974 (2007).

72. Mazel, T. et al. Direct observation of microtubule pushing by cortical dynein in living cells. Mol Biol Cell 25, 95–106 (2014).

73. Murrell, M., Oakes, P. W., Lenz, M. & Gardel, M. L. Forcing cells into shape: the mechanics of actomyosin contractility. Nat Rev Mol Cell Biol 16, 486–498 (2015).

74. Rossier, O. M. et al. Force generated by actomyosin contraction builds bridges between adhesive contacts. The EMBO journal 29, (2010).

75. Manneville, J.-B., Jehanno, M. & Etienne-Manneville, S. Dlg1 binds GKAP to control dynein association with microtubules, centrosome positioning, and cell polarity. Journal of Cell Biology 191, 585–598 (2010).

76. Etienne-Manneville, S. Microtubules in cell migration. Annu Rev Cell Dev Biol 29, 471–499 (2013).

77. Etienne-Manneville, S. & Hall, A. Integrin-mediated activation of Cdc42 controls cell polarity in migrating astrocytes through PKCzeta. Cell 106, 489–498 (2001).

78. Hashimoto-Tane, A. et al. Dynein-driven transport of T cell receptor microclusters regulates immune synapse formation and T cell activation. Immunity 34, 919–931 (2011).

79. Schnyder, T. et al. B Cell Receptor-Mediated Antigen Gathering Requires Ubiquitin Ligase Cbl and Adaptors Grb2 and Dok-3 to Recruit Dynein to the Signaling Microcluster. Immunity 34, 905–918 (2011).

