## Supplementary material Theory for "Cell adhesion and spreading on fluid membranes through microtubules-dependent mechanotransduction"

### Supplemental Material: theory

#### I. THEORETICAL MODEL AND DYNAMIC EQUATIONS

##### A. Model and free energy

Our theoretical model for the experiments consists of two membranes with height profiles $h_1(\mathbf{r})$ (SLB) and $h_2(\mathbf{r})$ (cell membrane), where $\mathbf{r}$ is the two-dimensional position vector and $h = h_2 - h_1 > 0$ is the inter-membrane separation. Ligands are bound to the SLB and the cell membrane is partially occupied by receptors. Pairs of ligands and receptors with the same $\mathbf{r}$ coordinate can form bonds. Otherwise, we refer to them as free. Bonds are connected to the cytoskeleton via a complex of adaptor proteins. We coarse grain over these complexes and refer to them simply as bonds. We denote the surface fraction of bonds as $\phi_b$, the surface fraction of free ligands in the SLB as $\phi_l$, and that of free receptors in the cell membrane as $\phi_r$. The setup is illustrated in Supp. Fig. S7 A.

In the absence of active cellular forces, the system can be described by the free energy (in units of $k_BT$)

$$F = \int \mathrm{d}^2 r \Bigg[ \phi_l \ln \phi_l + (1-\phi_b-\phi_l)\ln(1-\phi_b-\phi_l) + \phi_r \ln\phi_r + (1-\phi_b-\phi_r)\ln(1-\phi_b-\phi_r)$$
$$+ \phi_b \ln\phi_b - (1-\phi_b)\ln(1-\phi_b) - \epsilon\phi_b - \frac{1}{2}\chi\phi_b^2$$
$$+ \frac{1}{2}k_s h_1^2 + \frac{1}{2}k_0(h-l_0)^2 + \frac{1}{2}\phi_b k_b (h-l_b)^2 + \frac{1}{2}\sum_{j=1}^{2}\left(\kappa_j(\nabla^2 h_j)^2 + \sigma_j|\nabla h_j|^2\right)\Bigg]. \tag{1}$$

The first six terms account for the entropy of mixing of the free ligands, free receptors, and bonds, assuming that each occupies a comparable area (i.e. the lattice site). This entropy relies on the fact that the ligands are mobile, as is adequate on a fluid substrate, as opposed to a solid one. The last term $-(1-\phi_b)\ln(1-\phi_b)$ accounts for the fact that the bonds occupy both membranes simultaneously.

The term $-\epsilon\phi_b$ describes the total energy gain due to a binding energy $\epsilon > 0$ per bound ligand-receptor pair, while the term $-\frac{1}{2}\chi\phi_b^2$ describes short-range interactions between the bonds. A positive interaction parameter, $\chi > 0$, indicates attraction. Later, we will consider that the interaction depends on the force applied by the cytoskeleton, $f$.

The last line of Eq. (1) describes the free energy associated with changes in the membrane heights. The first term $\sim k_s h_1^2$ accounts for the short-range interaction between the SLB and the surface and wetting layer beneath it. We define the preferred height of the SLB to be $h_1 = 0$. The term proportional to $k_0(h-l_0)^2$ in the free energy accounts for the short-range interaction between the membranes (e.g., due to glycocalyx) that defines the preferred inter-membrane separation $l_0$ in the absence of bonds. In contrast, the bond is described as a harmonic spring with a spring constant $k_b$ and preferred length $l_b < l_0$. The last terms in Eq. (1) account for the spatial deformation of the membranes [1] in terms of the bending rigidities $\kappa_j$ and surface tensions $\sigma_j$.

Next, we apply the general approach of close-to-equilibrium, linear thermodynamics [2] to derive dynamic equations for the concentrations of the free ligands, free receptors, bonds, and the positions of the membranes. For simplicity, we consider only dissipative couplings between fluxes and their forces (described by a diagonal matrix of Onsager transport coefficients).

##### B. Dynamic equations

The concentrations of the free ligands , free receptors, and bonds are determined from the continuity equation (mass conservation) and binding and unbinding kinetics. The free ligands and bonds evolve according to

$$\frac{\partial\phi_l}{\partial t} = \frac{1}{\zeta_l}\nabla\left(\phi_l\nabla\frac{\delta F}{\delta\phi_l}\right) - k_{\mathrm{on}}\phi_{lr} + k_{\mathrm{off}}\,\phi_b, \tag{2}$$
$$\frac{\partial\phi_b}{\partial t} = \frac{1}{\zeta_b}\nabla\left(\phi_b\nabla\frac{\delta F}{\delta\phi_b}\right) + k_{\mathrm{on}}\phi_{lr} - k_{\mathrm{off}}\,\phi_b. \tag{3}$$

The first terms on the right-hand sides of Eqs. (2) and (3) correspond to two-dimensional diffusion within the membranes, written in terms of the ligand and bond friction coefficients ($\zeta_l$ and $\zeta_b$, respectively) that are inversely proportional to the diffusion coefficients. These diffusion terms vanish for immobile ligands on solid substrates. The last two terms describe the binding and unbinding kinetics, in terms of the binding rate $k_{\rm on}$, the surface fraction of pairs of free ligands and receptor, $\phi_{lr}$, and the unbinding rate $k_{\rm off}$. As the equilibrium surface fractions of free ligands and receptors in chemical contact with reservoirs are proportional to $1-\phi_b$ (see below), we have $\phi_{lr} = \phi_l\phi_r/\left(1-\phi_b\right)$. This is the equilibrium result of a microscopic lattice model that considers five microscopic states on each lattice site (that correspond to the occupancy states of both membranes): lipid-lipid, lipid-ligand, receptor-lipid, receptor-ligand, and bound receptor-ligand. The binding and unbinding rates depend on system parameters, such as the inter-membrane separation, $h$, and binding energy, $\epsilon$, as is detailed below.

The receptors follow similar dynamics with the addition that free receptors can be exchanged between the cell membrane and pools of receptors in the interior of the cell via endocytosis and exocytosis. Assuming reaction-limited kinetics [3, 4], the adsorption rate depends on the local chemical-potential difference with respect to the reservoir in the cytoplasm. Explicitly, the receptors evolve according to

$$\frac{\partial\phi_r}{\partial t} = \frac{1}{\zeta_r}\nabla\left(\phi_r\nabla\frac{\delta F}{\delta\phi_r}\right) + \frac{1}{\tau}\left(\mu_r^0 - \frac{\delta F}{\delta\phi_r}\right) - k_{\rm on}\phi_{lr} + k_{\rm off}\,\phi_b, \tag{4}$$

where $\zeta_r$ is the friction coefficient for the two-dimensional diffusion of free receptors within the cell membrane, $\tau$ is a characteristic adsorption time for molecules near the cell membrane, and $\mu_r^0$ is the receptor chemical potential in the cytoplasm.

The membrane dynamics are similarly determined from the derivatives of the free energy, according to

$$\frac{\partial h_1}{\partial t} = -\frac{1}{\Gamma}\frac{\delta F}{\delta h_1}, \tag{5}$$

$$\frac{\partial h_2}{\partial t} = -\frac{1}{\Gamma}\left(\frac{\delta F}{\delta h_2} - \phi_b f\right). \tag{6}$$

These equations describe the relaxation of forces exerted on the membrane due to deviations from their steady-state values. For simplicity, we have assumed the same dynamic coefficient $\Gamma$ for both the cell membrane and the SLB.

Importantly, the profile of the cell membrane is affected by an active, vertical force $\phi_b f$ that is exerted by the cytoskeleton on the bonds. In its absence, the system described above is in thermodynamic equilibrium. As the experiments suggest vertical microtubules-dependent forces, it is plausible that the force per bond, $f$, is fixed. This is in contrast to forces from larger complexes, where the total force is expected to be fixed, and the force per bond is determined by the number of bonds [5]. The exact dependence of $f$ on $\phi_b$ can be non-trivial and its determination requires further force measurements in adhesion sites of different bond densities.

Note that the dynamic Eqs. (2)-(6) are all within mean field (MF). Fluctuations in the number of bonds can be included via a one-step master equation [4, 6] and are important for determining the life time of clusters. Membrane fluctuations were also shown to be important in mediating effective cis-interactions in GUVs [7]. Such fluctuations can be incorporated in our theory by adding noise terms to Eqs. (5), (6).

##### C. Binding and unbinding rates and short-range interactions

The binding and unbinding rates ($k_{\rm on}$ and $k_{\rm off}$, respectively) determine the number of bonds at steady state according to $\phi_b/\phi_{lr} = k_{\rm on}/k_{\rm off}$ [Eq. (3)]. The rates depend on several of the model parameters. Binding requires overcoming an elastic stretching energy $\frac{1}{2}k_b\left(h-l_b\right)^2$ per bond when the inter-membrane separation, $h$, is different from the preferred length of the bonds, $l_b$, while unbinding requires overcoming an energy $\epsilon + \chi\phi_b$. Kramers' theory implies that the transition rates depend exponentially on these free-energy differences, such that

$$k_{\rm on} = k_0 \exp\left[-\frac{1}{2}k_b(h-l_b)^2\right], \tag{7}$$

$$k_{\rm off} = k_0 \exp\left[-\epsilon - \chi\phi_b + f/f_0\right], \tag{8}$$

where $k_0$ is an intrinsic timescale.

In Eq. (8) we have taken into account that the unbinding rate grows exponentially with the vertical force [5], where $f_0$ is a force scale. This can also be justified in the framework of Kramers' theory [8], where $f_0 = k_BT/x$ and $x$ is the change in the reaction coordinate (distance between the ligand and receptor) during unbinding. Assuming that the potential that binds the ligand and receptor together has a range of order of 0.5 nm, $f_0$ is on the order of 10 pN [9, 10]. At thermodynamic equilibrium ($f = 0$), the bond concentration that is obtained from the binding and unbinding rates minimizes the free energy, $\delta F/\delta\phi_b = 0$ (see next subsection).

The short-range attractive interaction between bonds involves mechano-sensitive adaptor proteins [11], such as talin, that exposes vinculin binding sites [12, 13] upon stretching. Applied forces change the protein conformation and lead to stronger effective attraction between bonds. As we coarse grain over the adaptor proteins, this effect is best accounted for phenomenologically via a force-dependent interaction parameter, $\chi(f)$. We consider the form

$$\chi(f) = \frac{\chi_0}{\left[1 + \exp\left(\Delta - f/\bar{f}\right)\right]^2}, \tag{9}$$

which can be justified by a microscopic two-state model that considers two bond states: a "native" state, and an "activated" one that has a typical attraction of order $\chi_0$. There is an energy barrier, $\Delta$, between the two states (in units of $k_BT$) and a force scale, $\bar{f}$, for activating the bonds. Considering transition rates that follow the same arguments as the binding and unbinding rates [Eqs. (7) and (8)], the concentration of activated bonds is expected to scale as $1/\left[1 + \exp\left(\Delta - f/\bar{f}\right)\right]$. Given typical conformation changes of the talin on the order of 30 nm [13], the force scale $\bar{f}$ is expected to be on the order of 0.1 pN.

##### D. Steady state solutions

We look for homogeneous steady states of Eqs. (2)-(6), which we denote by tildes. From Eq. (6), the solution for the membrane separation in terms of the bond density is

$$\tilde{h} = \frac{k_0 l_0 + \tilde{\phi}_b k_b l_b + f\tilde{\phi}_b}{k_0 + \tilde{\phi}_b k_b}. \tag{10}$$

Meanwhile, Eqs. (2) and (3) are simultaneously satisfied when $k_{\rm off}\phi_b = k_{\rm on}\phi_{lr} = k_{\rm on}\tilde{\phi}_l\tilde{\phi}_r/\left(1 - \tilde{\phi}_b\right)$. The receptor fraction is then found from the adsorption term in Eq. (4) as $\tilde{\phi}_r = \bar{\phi}_r\left(1 - \tilde{\phi}_b\right)$, where we have denoted the constant $\bar{\phi}_r = 1/\left(1 + e^{-\mu_r^0}\right)$. Finally, the steady-state concentration of the ligands is determined from a boundary condition. Given their mobility, we equate the free-ligand chemical potential with a reservoir. Similarly to the receptors, this yields $\tilde{\phi}_l = \bar{\phi}_l(1 - \tilde{\phi}_b)$.

Combining the results above for $\tilde{h}$, $\tilde{\phi}_r$ and $\tilde{\phi}_l$, together with the explicit expressions for the binding and unbinding rates [Eqs. (7) and (8)], leads to a transcendental equation for $\tilde{\phi}_b$,

$$\ln\left(\frac{\tilde{\phi}_b}{1 - \tilde{\phi}_b}\right) - \chi(f)\tilde{\phi}_b + \frac{1}{2}k_b(l_0 - l_b)^2\left(\frac{1 + c\tilde{\phi}_b f/f_b}{1 + c\tilde{\phi}_b}\right)^2 = \epsilon - \frac{f}{f_0} + \ln(\bar{\phi}_l\bar{\phi}_r). \tag{11}$$

Here, $c = k_b/k_0$ is the ratio between the spring constant of the bond and the inter-membrane interaction, and $f_b = k_b(l_0 - l_b)$ is the tension in a bond stretched to length $l_0$.

Eq. (11) corresponds to the condition that the chemical potential of bonds is constant. It can be interpreted as the binodal line of a binary mixture of bonds and vacancies in the mid-plane between the membranes. The right-hand side corresponds to the bulk chemical potential. The first term on the left-hand side originates in the entropy of mixing of bonds, as is allowed by the mobility of ligands and bonds. The second term accounts for the short-range attraction between bonds, as is mediated by adaptor proteins. It is stronger for larger forces, as is evident from Eq. (9).

The third term in Eq. (11) describes an effective membrane-mediated interaction at the MF level [14]. For small forces, $f < f_b$, bonds decrease the inter-membrane separation and the interaction is attractive. Bonds accumulate in order to bring the inter-membrane separation $h$ closer to the bond rest length $l_b$ and minimize the energy. For larger forces, $f > f_b$, vertical cytoskeletal forces separate the membranes further apart. Bonds then repel each other in order to bring $h$ closer to $l_b$.

Phase separation occurs when the effective attraction between bonds is strong compared with the mixing term. In this case, the left-hand side of Eq. (11) is non-monotonic in $\tilde{\phi}_b$ and the transcendental equation has multiple solutions. This is explained in detail below.

##### E. Condition for phase separation

To highlight the role of ligand affinity on clustering (or phase separation), we re-arrange Eq. (11) so as to separate the binding energy from the other terms,

$$\epsilon = \ln\left(\frac{\tilde{\phi}_b}{1-\tilde{\phi}_b}\right) - \chi(f)\tilde{\phi}_b + \frac{1}{2}k_b(l_0 - l_b)^2\left(\frac{1+c\tilde{\phi}_b f/f_b}{1+c\tilde{\phi}_b}\right)^2 + \frac{f}{f_0} - \ln(\bar{\phi}_l\bar{\phi}_r) \equiv G(\phi_b), \tag{12}$$

and we denote the right-hand side of this equation by a function $G$ of the bond concentration. When the mixing entropy term dominates, the function $G(\phi_b)$ is monotonically increasing, with $G \to -\infty$ as $\phi_b \to 0^+$ and $G \to +\infty$ as $\phi_b \to 1^-$. In such cases, there is a single well-mixed phase and the bond concentration increases with binding energy.

If the attractive interactions between bonds are strong enough, the chemical potential becomes non-monotonic, and the function $G(\phi_b)$ has two turning points, $0 < \phi_* < \phi_{**} < 1$ with $G(\phi_*) = G_{\max}$ being a local maximum of $G$ and $G(\phi_{**}) = G_{\min}$ being the local minimum of $G$. If the binding energy lies in the interval $G(\phi_{\min}) < \epsilon < G(\phi_{\max})$, then Eq. (12) admits three solutions. It is easy to check from the dynamic equations that the dilute solution, $\phi_1 < \phi_*$, and the dense solution, $\phi_3 > \phi_{**}$, are stable, while the middle solution $\phi_2$ is unstable.

When the binding energy lies in a favourable interval, $G_{\min} < \epsilon < G_{\max}$, and there exist two stable homogeneous steady states, the system can separate into a dilute and a dense phase, the latter of which appears by formation of clusters. For realistic parameter values, the upper bound $G_{\max}$ is on the order of 100 $k_BT$, much higher than the known binding energies for ligand-receptor pairs [15, 16]. Hence, we interpret the lower bound $\epsilon > G_{\min}$ as the condition for clustering, and we refer to $\epsilon_c = G_{\min}$ as the "critical binding energy".

#### II. RESULTS

We focus our theoretical predictions on two key aspects of the experiments on cell adhesion to fluid membranes: the role of receptor-ligand affinity via the binding energy, $\epsilon$, and the role of the cytoskeleton which is quantified by the vertical force, $f$, applied to each bond (e.g., by the microtubules and dyneins).

We consider a range of affinities from 5 to 15 $k_BT$, an interval which contains the binding energy of integrin-RGD at around 10 $k_BT$ [15]. Relative to this value, we estimate the binding energy of integrin-invasin to be around 15 $k_BT$ since the integrin-invasin pair can have a dissociation constant, $K_D \sim \exp(-\epsilon)$, about 200 times smaller than integrin-RGD [17]. We further consider a range of forces from 0 to 3 pN, which reasonably contains the typical forces applied to a single bond. Previous experiments on fluid substrates have found that individual integrin receptors apply forces smaller than 4.7 pN in integrin clusters outside of podosomes [18]. It has also been suggested that the connection between integrins and the cytoskeleton breaks under a force of about 2 pN [19]. Hence, the cytoskeletal force per bond in our model, which requires a viable connection between integrins and the cytoskeleton, is within the range we have considered.

Unless otherwise stated in the captions or legends of the figures, the theoretical results presented in this section are based on the default parameters $k_b = 0.25$ pN/nm [20], $k_0 = 0.01$ pN/nm [21], $l_0 = 100$ nm (typical size of glycocalyx), $l_b = 20$ nm (typical size of invasin), $\chi_0 = 4\ k_BT$ (order of magnitude estimate), $\Delta = 10\ k_BT$ [22], $f_0 = 10$ pN [9, 10], $\bar{f} = 0.1$ pN [23], $\bar{\phi}_r = 1\%$ and $\bar{\phi}_l = 8\%$ (estimated from the present experimental work).

##### A. Clustering as phase separation

We begin by describing the system through its phase diagram in the space of binding energies and cytoskeletal forces, in Supp. Fig. S7 B. As discussed at the end of the last section, the binding energy must reach a critical value (white line on the phase diagram) in order for the system to stably sustain two homogeneous steady states: a dense phase and a dilute phase (in the experiments, integrin clusters and background concentration). A striking feature of the phase diagram is that the critical binding energy decreases at intermediate forces. This feature is due to the adaptor-protein mediated attraction between bonds, and colorblueis discussed again below. In early nascent adhesions, the cytoskeletal forces are weak and this effect is not yet apparent. We hence focus on the small force regime and plot the bond concentration against the binding energy in Supp. Fig. S7 C. Our model predicts that the dense phase (clusters) appears through a saddle-node bifurcation (or first-order phase transition) at a critical binding energy which depends on the applied force, and thereafter the concentration of bonds inside the dense phase increases towards full surface coverage ($\phi_b \to 1$). If we increased the binding energy even further, we would observe that the unstable solution (dashed line) merges with the dilute stable solution in another saddle-node bifurcation, after which point only the dense phase survives. However, this goes beyond the realistic range of receptor-ligand binding energies [15, 16].

##### B. Clustering under force

Next, we turn our attention to the vertical force applied to each bond by the cytoskeleton. Our model predicts that the clusters are able to sustain larger forces, and hence grow larger over time, if either the affinity of ligand-receptor pairs (Supp. Fig. S7 D) or the strength of attractive interactions through the adaptor proteins (Supp. Fig. S7 E) is increased. A non-trivial prediction of our model is that, for a specific range of binding energies (for example, $\epsilon = 9\ k_B T$ in Supp. Fig. S7 D) the clusters exist at very low forces and then again at slightly higher forces, with a gap in between. This disruption is due to the fact that the membrane-mediated attraction (which always decreases with increasing force, see third term in Eq. (11)) weakens before the attraction through the adaptor proteins has had a chance to kick in. In between, neither of the two attractive mechanisms is strong enough to compete with the mixing entropy of bonds, so the system is well-mixed.

##### C. Variation of parameters

Finally, we perform a systematic variation of the parameters to highlight the effect of a few key quantities on the conditions for clustering – Supp. Fig. S7 (F-J). As previously mentioned, the non-monotonic dependence of the critical binding energy on the applied force is due to the attraction mediated by adaptor proteins. As we increase the strength of this interaction, in Supp. Fig. S7 F, the critical binding energy is lowered, which is consistent with our understanding that adaptor proteins contribute to clustering. The effect only occurs above a threshold force ($f > \Delta\bar{f}$) which can be understood, for example, as the force required to stretch the talin and activate the bonds. In Supp. Fig. S7 G, we directly increase this threshold and observe, as expected, that the onset of the attractive mechanism mediated by the adaptor proteins is delayed. We also observe that increasing the harmonic spring constant of the bonds (Supp. Fig. S7 H), decreasing the strength of short-range interactions between the membranes (Supp. Fig. S7 I) and decreasing the preferred inter-membrane separation (Supp. Fig. S7 J) all have the effect of lowering the barrier for clustering. This is consistent with our understanding that, in order to form clusters, the bonds must be able to impose their preferred inter-membrane separation over the one imposed by the glycocalyx (hence, higher $k_b$ and lower $k_0$ favours clustering), and that this is more easily done when the difference between the two, $l_0 - l_b$, is lower. Note that we have also systematically varied the other parameters in our model (results not shown) and found no significant effect within the range of values relevant for the experiments.

#### III. CONCLUSION

Our theory predicts that ligand-receptor pairs with higher affinity lead to denser phases (Supp. Fig. S7 C) and are able to sustain larger vertical forces (Supp. Fig. S7 D). This is consistent with the experimental observation that clusters form with higher density on fluid membranes that contain high-affinity ligands such as invasin, as compared to RGD.

Our theory further predicts a non-trivial dependence of adhesion-cluster maturation on the force per bond, which we have considered here to be fixed. Small forces increase the effective attraction between bonds due to conformation changes in adaptor proteins. Larger forces per bond on the order of few pN possibly decrease the attraction via the membrane-mediated interaction, while even larger forces per bond on the order of 10 pN are expected to break bonds and suppress the formation of adhesion sites.

Finally, our theoretical predictions are consistent with the experimental observation that cluster maturation is accompanied by the recruitment of adaptor proteins, since clusters can sustain larger forces per bond when the attractive mechanism mediated by adaptor proteins is stronger (Supp. Fig. S7 E).

---

[1] S. A. Safran, *Statistical thermodynamics of surfaces, interfaces, and membranes* (CRC Press, 2018).
[2] S. R. De Groot and P. Mazur, *Non-equilibrium thermodynamics* (Courier Corporation, 2013).
[3] H. Diamant and D. Andelman, Kinetics of surfactant adsorption at fluid- fluid interfaces, The Journal of Physical Chemistry **100**, 13732 (1996).

[4] U. S. Schwarz and S. A. Safran, Physics of adherent cells, Reviews of Modern Physics **85**, 1327 (2013).

[5] G. I. Bell, Models for the specific adhesion of cells to cells: a theoretical framework for adhesion mediated by reversible bonds between cell surface molecules., Science **200**, 618 (1978).

[6] T. Erdmann and U. S. Schwarz, Stability of adhesion clusters under constant force, Physical Review Letters **92**, 108102 (2004).

[7] S. F. Fenz, T. Bihr, D. Schmidt, R. Merkel, U. Seifert, K. Sengupta, and A.-S. Smith, Membrane fluctuations mediate lateral interaction between cadherin bonds, Nature physics **13**, 906 (2017).

[8] R. Elber, eng*Molecular kinetics in condensed phases : theory, simulation, and analysis / Ron Elber, Dmitrii E Makarov, Henri Orland.* (2020).

[9] O. Thoumine, P. Kocian, A. Kottelat, and J.-J. Meister, Short-term binding of fibroblasts to fibronectin: optical tweezers experiments and probabilistic analysis, European Biophysics Journal **29**, 398–408 (2000).

[10] F. Li, S. D. Redick, H. P. Erickson, and V. T. Moy, Force measurements of the 51 integrin–fibronectin interaction, Biophysical Journal **84**, 1252 (2003).

[11] B. Geiger and A. Bershadsky, Exploring the neighborhood: adhesion-coupled cell mechanosensors, Cell **110**, 139 (2002).

[12] A. Del Rio, R. Perez-Jimenez, R. Liu, P. Roca-Cusachs, J. M. Fernandez, and M. P. Sheetz, Stretching single talin rod molecules activates vinculin binding, Science **323**, 638 (2009).

[13] M. Yao, B. T. Goult, B. Klapholz, X. Hu, C. P. Toseland, Y. Guo, P. Cong, M. P. Sheetz, and J. Yan, The mechanical response of talin, Nature communications **7**, 11966 (2016).

[14] S. Komura and D. Andelman, Adhesion-induced lateral phase separation in membranes, The European Physical Journal E **3**, 259 (2000).

[15] S. Goennenwein, M. Tanaka, B. Hu, L. Moroder, and E. Sackmann, Functional incorporation of integrins into solid supported membranes on ultrathin films of cellulose: Impact on adhesion, Biophysical Journal **85**, 646 (2003).

[16] M. Wilchek and E. A. Bayer, Introduction to avidin-biotin technology, in *Avidin-Biotin Technology*, Methods in Enzymology, Vol. 184, edited by M. Wilchek and E. A. Bayer (Academic Press, 1990) pp. 5–13.

[17] G. Van Nhieu and R. Isberg, The *Yersinia pseudotuberculosis* invasin protein and human fibronectin bind to mutually exclusive sites on the $\alpha_5\beta_1$ integrin receptor, Journal of Biological Chemistry **266**, 24367 (1991).

[18] R. Glazier, J. M. Brockman, E. Bartle, A. L. Mattheyses, O. Destaing, and K. Salaita, DNA mechanotechnology reveals that integrin receptors apply pN forces in podosomes on fluid substrates, Nature Communications **10**, 1 (2019).

[19] G. Jiang, G. Giannone, D. R. Critchley, E. Fukumoto, and M. P. Sheetz, Two-piconewton slip bond between fibronectin and the cytoskeleton depends on talin, Nature **424**, 334 (2003).

[20] H. Gao, J. Qian, and B. Chen, Probing mechanical principles of focal contacts in cell-matrix adhesion with a coupled stochastic-elastic modelling framework, Journal of the Royal Society Interface **8**, 1217 (2011).

[21] The estimate for $k_0$ is based on a typical membrane fluctuation $\sqrt{\langle h^2 \rangle}$ of 30 nm (measured for GUVs by Fenz *et al.* [7]), with $k_0 = 2k_BT/\langle h^2 \rangle$.

[22] Considering a typical conformation change of the talin on the order of 30 nm [13] and the fact that a force of 5 pN is sufficient to uncover the first vinculin binding sites [24], we expect $\Delta$ to be smaller than 5 pN $\times$ 30 nm = 36 $k_BT$.

[23] The estimate for $\bar{f}$ is based on a typical conformation change of the talin on the order of $\bar{l} = 30$ nm [13], with $\bar{f} = k_BT/\bar{l}$.

[24] J. Yan, M. Yao, B. T. Goult, and M. P. Sheetz, Talin Dependent Mechanosensitivity of Cell Focal Adhesions, Cellular and Molecular Bioengineering **8**, 151 (2015).
