## Supplementary material video annotations for "Cell adhesion and spreading on fluid membranes through microtubules-dependent mechanotransduction"

Supplementary video files annotations.

Supplementary video V1 :

Brightfield time-lapse imaging of a trembling MEF cell on Invasin-SLB. Frames were acquired at an interval of 10 sec and video playback speed is at 7 fps. Time in hh:mm. Scale bar: 5 µm.

Supplementary video V2 :

Brightfield time-lapse imaging of an adherent MEF cell on Invasin-SLB. Frames were acquired at an interval of 10 sec and video playback speed is at 7 fps. Time in hh:mm. Scale bar: 5 µm.
