## Supplementary material for "Cell adhesion and spreading on fluid membranes through microtubules-dependent mechanotransduction": Materials and Methods

**Cell chamber assembly**

For the bilayer to be always hydrated it is formed in a sample flow chamber that always has buffer inside. The sample chamber for the cell adhesion experiment is a custom-made flow chamber that consists of a 26 mm x 76 mm coverglass slide and a 1.5 thickness coverslip from VWR glued together by two melted stripes of parafilm. Prior to chamber assembly the coverslip is cleaned and activated in a series of sonication steps of 30 min in the water bath sonicator (Elmasonic S 10 (H)): in distilled water, 2% Hellmanex III, distilled water, 1M KOH, distilled water. The KOH has a dual role of cleaning and glass activation. At the end of the cleaning steps the coverslip is dried under nitrogen flow and assembled to form the microscopy chamber. Coverglass slides are simply rinsed with EtOH, H_2_0, EtOH and blow dried with nitrogen.

The distance between the two glass slides (the height of the channel) is about 250 µm. The chamber volume is about 50 µl. The assembled chambers can be stored in closed Petri dishes (protected from dust) at room temperature for about 3 days.

**Small Unilamellar Vesicles (SUVs) preparation**

1,2-dioleoyl-*sn*-glycero-3-phosphocholine (DOPC), 1,2-dioleoyl-sn-glycero-3-[(N-(5-amino-1-carboxypentyl)iminodiacetic acid)succinyl] nickel salt (DGS-NTA(Ni)), 1,2-dioleoyl-sn-glycero-3-phosphoethanolamine-N-[4-(p-(cysarginylglycylaspartate-maleimidomethyl)cyclohexane-carboxamide] sodium salt (DOPE-RGD) were purchased from Avanti Polar Lipids (Alabaster, AL, USA). Invitrogen™ Marina Blue™ 1,2-dihexadecanoyl-*sn*-glycero-3-phosphoethanolamine (Marina Blue™ DHPE) was purchased from Invitrogen (Waltham, MA, USA). SLBs were formed by fusion of Small Unilamellar Vesicles (SUVs) on the coverslip. We used two lipid compositions to prepare SUVs and consequently SLBs: 1) DOPC/DGS-NTA(Ni)/DHPE-Marina Blue (94/2/4, mol/mol); 2) DOPC/DOPE-RGD/DHPE-Marina Blue (94/2/4). Similar lipid compositions were successfully used by other groups to prepare SLBs^1^. The SUVs were prepared using the following protocol:

1. Lipids solubilized in chloroform are mixed together in a glass vial at 1 mg/ml, blow dried with a nitrogen flow, placed in vacuum desiccator for 1 hour, then rehydrated with distilled water for 15 min at room temperature, to a final lipid concentration of 1 mg/ml;
2. After rehydration, the glass vial is vortexed to detach the liposomes;
3. The solution is sonicated for 30 min in the water bath sonicator (Elmasonic S 10 (H));
4. The solution is then centrifuged at 20k RCF for 1 hour;
5. The solution is filtered through a 200 nm filter (Millipore) with a syringe;
6. The sample is dissolved in the SUV fusion buffer at the final composition: 10 mM Tris pH 7.3; 120 mM NaCl.

Lipids preparations are closed under with argon at all stages of SUV preparation to minimize their degradation by oxidation. The prepared SUVs were not stored but used immediately to prepare SLBs.

**Invasin preparation**

While RGD-ligands bind to a larger repertoire of integrins (α_5_β_1_, α_8_β_1_, α_V_β_1_, α_V_β_3_, α_V_β_5_, α_V_β_6_, α_V_β_8_ and α_II_β_3_)^2^, Invasin binds to a subset of β_1_-integrins (α_3_β_1_, α_4_β_1_, α_5_β_1_ and α_6_β_1_), including the fibronectin integrin α_5_β_1_, with which Invasin has a dissociation constant (K_d_) two orders of magnitude lower than that of RGD peptide^3^.

- **Plasmid preparation**

The plasmid for Invasin expression was constructed in two steps. First, a DNA sequence coding for a TEV cleavage site followed by a 6xHis tag followed by the sequence coding for the last 474 amino acids of Invasin, inv474 (PDB: 1CWV) (in 5 3 direction) and flanked by SacI and HindIII restriction sites was synthetized and subcloned by Thermo Fisher Scientific. Second, the insert of interest was cloned into a pMal p5x expression vector by digesting with SacI and HindIII enzymes and ligation. As a result, we have obtained the plasmid that expresses a periplasmic maltose binding protein (MBP) fused to Invasin in *E.Coli* periplasm under IPTG inducible P_tac_ promoter (pOM3474).

- **Invasin expression**

For Invasin expression we followed the previously described protocol^4^ and expressed Invasin in the *E. Coli* periplasm to insure the oxidizing environment for the proper disulfide bond formation. We used an *E. Coli* strain lacking *DegP* protease^4^. Proteins were expressed in 2YT medium + selection antibiotics (ampicillin for the plasmid and kanamycin for the *E. Coli* strain at final concentrations 100 µg/ml and 50 µg/ml, respectively) + 0.2% glucose. Glucose is needed to inhibit amylase expression that can later perturb MBP-Invasin purification. Bacterial culture was inoculated from a single colony and incubated at 30 °C while shaking until the OD_600_ = 0.5−0.6. Then cells were induced with 0.5 mM IPTG for 4 hours. MBP-Invasin is well expressed at the right molecular size (approximately 96 kDa), thus we can proceed with the protein production, purification and labelling at the bigger scale.

- **Invasin purification**

Invasin purification and labelling was performed in several steps:

1. Affinity purification using an amylose resin (New England Biolabs);

2. MBP cleavage by TEV protease;

3. Protein labeling with a succinimidyl ester (SE, Tocris) reactive dye (optional);

4. Purification by size exclusion chromatography using a Superdex 75 column.

The more detailed protocol illustrates the purification process from 2 liters of bacterial culture that expressed MBP-Invasin:

**Cell lysis**

1. Resuspend cell pellets in approximately 80 mL Lysis Buffer: 25 mM HEPES pH 7.3, 500 mM NaCl, 1 mM EDTA;

2. Add a final of 1x EDTA Free cOmplete protease inhibitor tablet (Roche), 1 mM PMSF, 10 µg/ml DNase and 10 µg/ml lysozyme;

3. Cells were lysed via sonication (misonix sonicator ultrasonic processor XL, large probe at 20% intensity) on ice at 70% intensity: 3 sec - On, 3 Sec Off for 5 minutes;

4. Sample was centrifuged at 20,000 RCF for 60 Min, 4°C and supernatants collected; Affinity purification against the MBP;

5. 10 mL Amylose Resin were pre-washed in water then Lysis buffer;

6. Cleared extracts were applied to Amylose Resin, and allowed to bind using a tube roller for 2h, 4°C;

7. Samples were washed 2x 40mL using batch method with Wash Buffer 1 (+EDTA);

8. Amylose beads were applied to a column and washed a further 10x CV with the same buffer;

9. Proteins were eluted in 15 mL of Elution Buffer: 25 mM HEPES pH7.3, 500 mM NaCl, 10 mM maltose (the steps of the affinity purification against the MBP were repeated);

10. Protein concentration was estimated by measuring the absorbance spectrum at the Cary Eclipse Fluorescence spectrophotometer; the samples were analyzed by protein electrophoresis on SDS-PAGE gels.

**TEV cleavage and labelling with JF549 SE dye**

11. Add TEV protease at about 1:100 (TEV/Invasin, mol/mol);

12. Add JF549 SE dye (Tocris) at about 1:2 (Invasin/dye, mol/mol);

13. Incubate at 4°C in aluminum foil on the rolling table o/n; Size exclusion purification;

14. Labelled samples were concentrated down to approximately 500 µL and purified further by size exclusion chromatography using a Superdex 75 column. Buffer: 25 mM HEPES pH 7.3, 120 mM NaCl. The elution fractions were analyzed by protein electrophoresis on SDS-PAGE gels.

The fractions corresponding to Invasin were collected and its absorbance spectrum was by measured at the Cary Eclipse Fluorescence spectrophotometer. Invasin concentration C and labelling ratio n^∗^ (the labeling molar ratio of the fluorescence species) were estimated from the sample absorbance measurements A_280_ and A_max_ (absorbances at λ = 280 nm and λ corresponding to the fluorescence of the dye which for the JF549 is 549 nm), the extinction coefficients of JF549 dye ϵ_dye_ (from tocris.com) and Invasin ϵ_prot_ (expasy.org/protparam) 101000 [M^−1^ cm^−1^] and 57995 [M^−1^ cm^−1^] , respectively, and the CF correction coefficient that is equal to 0.169 (from tocris.com):

C = (A_280_ − A_max_ · CF)/ ϵ_dye_;

n^∗^ = A_280_/ ϵ_prot_ · C.

**Supported lipid bilayer (SLB) preparation**

The protocol is inspired by the protocol developed in the group of Jay Groves^1,5^.

The SUV solution is incubated in the sample chamber for 30 min to allow the SUV fusion on the substrate. Then the unfused vesicles are washed away with 10 x chamber volumes of the SUV fusion buffer. Next, the SLBs are washed with 10 x chamber volumes of the cell buffer: 25 mM HEPES pH 7.3; 120 mM NaCl; 7 mM KCl; 1.8 mM CaCl_2_; 0.8 mM MgCl_2_; 5 mM glucose. Any remaining defect on the surface is then passivated by incubation of the chamber in a "blocking" solution made of the cell buffer complemented with 0.1 mg/ml β-casein for 15 min. Then the SLBs is washed again with 10 x chamber volumes of the cell buffer to remove the excess of the blocking solution.

To prepare Invasin-coated SLBs, bilayers containing Ni-lipids are functionalized with 6xHis-tagged Invasin following the protocol^1^. SLBs are incubated with 400 nM Invasin in the cell buffer for 1 hour at room temperature. Then the unbound proteins are washed away with 10 x chamber volumes of the cell buffer in two steps spaced in time by 30 min.

**SLB quality and fluidity control**

We prepared RGD-SLB and SLB(Ni), functionalized SLB(Ni) with fluorescently labelled Invasin or EGFP (as a control). To assess the quality of SLBs (absence of defaults) they were inspected visually under the microscope. We observed fluorophore distributions in both DHPE-Marina-Blue and fluorescent proteins. If no defaults were detected, we proceeded further and tested the SLBs on fluidity. The fluidity of the prepared SLBs were checked by using the fluorescence recovery after photobleaching (FRAP) technique (Figure S1A). We can deduce the diffusion coefficients of fluorescent lipids or proteins attached to the bilayer by analyzing the fluorescence recovery curves after photobleaching of a small area of the SLB^6^. For each sample we have measured 10 FRAP curves in 2 experiments from a photobleached circular zone of approximately 20 µm^2^. A typical normalized fluorescence recovery curves for lipids (DHPE-Marina Blue) of the SLB(Ni) in presence of Invasin and Invasin on the SLB(Ni) are shown in the Figure S1B. They were fitted with the following function:

f(t) = f_0_ + (f_plateau_ − f_0_)(1 − e^−kt^),

from which we can find the immobile fraction of fluorophores as 1 – f_plateau_. The interaction of the lipid bilayer with the solid surface is limited since the immobile fraction (Figure S1C) for fluorescent lipids is lower than 7% for SLB-RGD, and even lower for SLB-Ni. Lipids remain mobile when Invasin or EGFP is bound to the SLB-Ni. We also see that the immobile fraction of Invasin on the SLB is of the order of 11%, and 5% for EGFP. The diffusion coefficient D, the radius of the bleached area r and the half-recovery time t_1/2_ are link by the following expression: D = 0.224 · r^2^/t_1/2_.

The diffusion coefficients for lipids are approximately 75% lower in SLB(Ni) than in SLB(RGD), maybe due to electrostatic interactions between the glass and the Ni lipids (0.8 versus 1.4 µm^2^/s, respectively) (Figure S1D). These values are in fairly good agreement with the previous studies of lipid diffusion in lipid bilayer (SLBs or liposomes)^7,8^. Binding of Invasin or EGFP to the bilayer does not change the lipid diffusion coefficient, showing that the bilayer is fluid. As expected, since proteins are bigger than lipids, the protein diffusion coefficients are lower, more than two times lower than that of the lipids (0.4 and 0.3 µm^2^/s for Invasin and EGFP respectively). These results show that the RGD-lipids and Invasin-lipids complexes in our experiments are mobile in the bilayer.

**Fluorescence calibration of SLBs**

The aim of the calibration is to quantify the density of fluorescence species in the cell adhesion plane by using supported lipid bilayers as fluorescence standards. We adapted the original protocol^5^ to perform the calibration. Within a certain range of fluorophore densities, we can relate the density of the fluorescence species n_fl_ [µm^−2^] and the intensity I_fl_ [AU] of the fluorescence image of the cell surface in the adhesive contact by the following:

N_fl_ = A_fl_ · I_fl_;

where A_fl_ is the proportionality factor that depends on the fluorescent molecule and imaging conditions. This linear relationship holds for concentrations below the critical concentration at which fluorophores start to self-quench^9^. The range of densities of fluorescence species where this linear relationship holds is verified during the calibration process.

Standards have to be used to calibrate the fluorescence in membranes, in illumination conditions similar to the experiments. In practice, fluorescent lipids are convenient since they can be incorporated in the bilayer at controlled concentration, thus their density is known and the corresponding fluorescence can be measured as a function of their density. Here, SLBs are particularly suitable if the fluorescence intensity is measured in the exact same conditions as in the experiments where the contact zone between cells and SLBs are imaged. But, to account for differences in fluorescence yield between the fluorescent markers on the lipid and the protein, the fluorescence of the lipids and the proteins at known concentrations must be compared; for this, measurements in bulk are well-suited.

The proportionality factor A_fl_ is related to the proportionality factor of the calibration standard A_st_ by the following relation:

A_fl_ = A_st_/ (F · n^∗^);

where the correction factor F = I_fl_/I_st_ is the ratio between the intensities of the fluorescence species I_fl_ and the standard I_st_ at a given concentration in solution and n^∗^ is the labeling molar ratio of the fluorescence species. n^∗^ = 1 for lipids, but might be not equal to 1 for proteins, depending on the efficiency of the labeling protocol. n^∗^ must be measured for each preparation of proteins with a spectrophotometer.

F considers the optical properties of the microscope and spectral differences between the fluorophore and the standard. In order to determine A_fl_ for a given fluorophore, we need to do the following:

1. Calibrate our imaging system with a standard fluorophore and measure the proportionality factor A_st_ in SLB with fluorescent lipids;

2. Measure the correction factor F with experiments in solution.

More precisely, in the first step, we prepare a series of SLBs with fluorescence lipids (standard) incorporated at known densities. Then we image these SLBs with exactly the same imaging conditions (laser powers, exposure times and fluorescence channels, or sets of fluorescence filters) as we do for the fluorophore of interest. We have taken at least 10 images in different areas of the SLB for every density of every studied fluorophore. In the second step, we measure the correction factor F, the dimensionless factor that represents the efficiency of the fluorophore versus the standard. Practically, to calculate F, we measure the ratio of intensities between the fluorophore and the standard at a given concentration in solution (in bulk), or more precisely, the ratio of the slope of the plots Intensity versus bulk concentration, for both fluorophores, α_st_ and β_fl_. This measurement must be done directly at the microscope with exactly the same imaging conditions as for the first step. We did 2 calibrations (in green and red channel) for the following molecules: AlexaFluor488 and JF549.

We used Bodipy FL DHPE (Molecular probes, referred to as BodipyFL in the following) and Texas Red DHPE (Invitrogen, referred to as TexasRed in the following) lipidated dyes as standards for the green and the red calibrations, respectively.

**Fluorophore preparation for calibration**

All fluorophores, except the lipidated ones, were diluted in the working cell buffer (25 mM HEPES pH 7.3; 120 mM NaCl; 7 mM KCl; 1.8 mM CaCl_2_; 0.8 mM MgCl_2_; 5 mM glucose). As lipids would aggregate in aqueous solution, we have solubilized them in detergent. Lipid solutions of BodipyFL or TexasRed were blow-dried under argon and vacuum for 30 minutes, and re-solubilized in the working cell buffer with 2.25 mM n-Dodecyl β-D-maltoside (DDM, Sigma Aldrich).

The accuracy of the calibration greatly relies on the precise knowledge of the fluorophore concentrations. The concentrations of all samples were checked on Cary Eclipse Fluorescence Spectrophotometer using the following published extinction coefficients: AlexaFluor488 (73000 [M^-1^cm^-1^]), BodipyFL (80000 [M^-1^cm^-1^]), JF549 (101000 [M^-1^cm^-1^]), TexasRed (116000 [M^-1^cm^-1^]), mCherry (72000 [M^-1^cm^-1^]).

The images of the fluorophores in bulk were taken approximately 10 µm above the coverslip. We observed that the intensity distribution of the fluorescence signal of a dye in solution is not homogenous in the imaging plane (Supplementary Fig. S3). This is related to the non-homogeneous illumination. The illumination seems to be more or less homogeneous in the middle of the illumination pattern “the plateau” (Supplementary Fig. S3A). For our analysis we either considered the intensity in this region or corrected the inhomogeneity of the illumination by using the fluorescence signal of bilayers. We have taken at least 10 images in different areas of the chamber for every concentration of every studied fluorophore.

The standard SLBs (DOPC + BodypyFL) with different concentrations of BodipyFL were prepared following the previously described protocol (“SLB preparation” in the Methods). We calculated the surface densities of BodipyFL in the SLBs from their molar ratios and assuming the area of a DOPC lipid projected on the SLB plane to be 0.72 nm^2^ and, thus, the number of lipids per µm^2^ to be 2 · µm^2^ / 0.72 nm^2^ = 2.8 · 10^6^ (where the factor 2 is to take both bilayer leaflets into account)^10^. For each sample 30 images were taken and the intensities of the plateau region were averaged. We plotted the density-intensity standard curves for both calibrations and also intensity-concentration curves for all fluorophores in bulk (Supplementary Fig. S3). Therefore, we obtained the following calibration values for our fluorescence species:

β_1_-integrin labelled with Alexa Fluor 488 (A_st_ = 3.146 µm^-2^; α_fl_ = 3410 µM^-1^; β_st_ = 1872 µM^-1^; F = 1.82; n* = 1; A_fl_ = 1.73 µm^-2^);

JF549 labelled Invasin (A_st_ = 3.857 µm^-2^; α_fl_ = 2366 µM^-1^; β_st_ = 1167 µM^-1^; F = 2.03; n* = 0.427; A_fl_ =4.45 µm^-2^).

Note that the calibration factors were measured from fluorescent dyes in solution and not from fluorescent proteins. F was shown to be about 3 times higher with AlexaFluor488 bound to anti-biotin than to streptavidin^5^. But, weaker differences were observed in other cases. In practice, we should purify the proteins of interest, label them and make the calibration, which is not possible. In addition, it is possible that the dye fluorescence is different in the cell environment as compared to the buffer. Thus, the absolute values of the protein densities in the adhesion structures that we deduce from our experiments might be off by some systematic factor, however, it cannot be by an order of magnitude.

According to the fluorescence calibration we found that Invasin was distributed homogeneously on SLBs and its density corresponded to approximately 600 Invasin/µm^2^ (under our SLB preparation protocol).

**Microscopy**

We have used a spinning disk microscope to image fluorescent lipid bilayers and cell adhesion on them. The microscope consists of a CSU-X1 Yokogawa head mounted on an inverted Ti-E Nikon microscope with a motorized XY stage (MadCity Lab®). Images were acquired with Metamorph software (Molecular Devices®) through a 100x NA1.45 objective with a Photometrics 95B-sCMOS camera. Live cell adhesion imaging was performed at 30°C using the stage top incubator (Tokai hit®). Fixed cells were imaged at RT with the same microscope. The setup is equipped with 4 Cobolt lasers from Hübner Photonics: 405 nm (100mW), 488 nm (100mW), 561 nm (50mW) and 633 nm (100mW). They allow sample imaging in 4 fluorescence channels (405, GFP, Cy3 and Cy5). Samples can also be imaged in wide field mode using the transmission light from LED source. The setup is also equipped with a FRAP photoactivation module.

We have used the same imaging conditions in our microscopy experiments: 405 (laser power: 15%; exposure time: 100ms), GFP (laser power: 30%; exposure time: 300ms), Cy3 (laser power: 30%; exposure time: 300ms) and Cy5 (laser power: 30%; exposure time: 300ms).

For imaging cells above the bilayer Z-stack images of 3 µm height are taken. The stack is centered at the SLB plane and its step is 0.3 µm.

Diffraction limit (DL), or optical lateral (xy plane) resolution was calculated as follows:

Res_xy_ = 0.51 · λ_em_ /NA,

Where λ_em_ – emission wavelength of a fluorophore, NA – numerical aperture of the objective.

**Image analysis**

- **“trembling/adherent” cells**

Cells were qualitatively classified as “trembling” if their edges but not their centers of mass moved at the time scale of 5 seconds. The edge movement was detected manually based on the bright field images.

- **Cell contour detection and measurement of cell morphology parameters**

Cell contours were manually detected from bright field images by using a polygonal selection tool in ImageJ. Then the “cell contour” selection was used to calculate the projected cell area A (“projected area”) and the cell circularity index C (“circularity”) as follows:

C = 4pi*A/P^2^, where P is the perimeter of the “cell contour” selection.

- **Integrin cluster detection and quantification**

Before detecting integrin clusters, we made a correction on the illumination inhomogeneity. The illumination is not homogeneous across the image (Supplementary Fig. S3A). Therefore, we introduced the "illumination" map to correct this issue. It consists in normalizing the image by that of a fluorescent supported lipid bilayer (SLB). The SLB has a homogeneous distribution of fluorescent lipids in the image plane, so it is a perfect candidate for the illumination map. The intensities of the illumination map range between 0 (the dimmest illumination) and 1 (the brightest illumination). Therefore, the image correction for the illumination inhomogeneity is obtained by the division of pixels intensities of the image by the intensities of the corresponding pixels of the illumination map.

To detect β_1_-integrin clusters we first used previously described fluorescence calibration transforming raw intensity images of integrins (corrected for illumination inhomogeneity) to integrin concentration maps. Second, we segmented these concentration maps by defining an integrin density threshold that separates two distributions (clusters and background) on the density histogram. The integrin density thresholds could be set manually or automatically by using an algorithm based on Renyi’s entropy thresholding. This thresholding method was previously described^11^ and is now one of the standard threshold methods available in ImageJ. The method defines a threshold intensity value that maximizes the informational entropy and entropic correlation of “cluster” and “background” distributions^12^.

- **Membrane tube detection**

To detect membrane tubes associated with β_1_-integrin clusters we first use “Reslice” function of ImageJ at the SLB channel to create cross sections perpendicular to the SLB and centered at integrin clusters. Next, we segment the images of the cross sections using Renyi’s entropy thresholding algorithm in order to find membrane tethers. In the segmented image we define a tube as an object larger than 1.5 µm in z dimension.

Custom-written codes used to analyze the data in the current study are available from the corresponding authors on reasonable request.

- **Localization of integrin clusters and membrane tubes in the adhesion plane of the cell**

To locate integrin clusters or membrane tubes in the adhesion plane of the cell with respect to the cell center and periphery we have defined the “cell center” and the “cell periphery” zones. The border between the two zones is resulted from a scaling transformation of the cell border, such that the cell periphery corresponds to the band parallel to the cell border and the cell center corresponds to the part of the cell that excludes the cell periphery. The two zones are chosen such that they have the same area.

Distances from integrin clusters to the cell border are defined as the shortest distance from the edge of the cluster to the cell border. Weighted mean of the distance between integrin clusters and the cell border is defined as the average distance of all detected integrin clusters in the cell weighted by the number of integrins in the clusters. For each cell, the weighted mean distance is normalized by the average distance of uniformly distributed clusters from the border of that particular cell, which is calculated numerically by discretizing the cell area into a fine rectangular mesh, placing one cluster at each grid point, and calculating the average distance of such uniformly distributed clusters to the cell edge. The mesh is iteratively refined until the error falls below 1% in a self-convergence test.

**FA protein, actin and microtubule enrichments in β_1_-integrin clusters**

Signal enrichment of the co-expressed fluorescent fusion proteins (FPs) for FA proteins, F-actin and microtubule imaging at β_1_-integrin clusters was calculated as follows. Mean fluorescence intensity of the FP was detected in the region of β_1_-integrin clusters and then was normalized by the mean intensity of the FP in the cell. It was compared with “shuffled control enrichments”, enrichment calculated at random pixels instead of β_1_-integrin clusters regions.

FA protein recruitment to integrin clusters experiments were done in fixed cells. Fluorescence calibrations were not performed in these conditions. β_1_-integrin clusters were defined using the same threshold (100 AU).

β_1_-integrin clusters were defined using the threshold of 100 integrins/µm^2^ for experiments of F-actin and microtubule recruitment to integrin clusters.

**Cell culture**

We used the following cell lines in our experiments:

Mouse Embryonic fibroblasts (MEF) β_1_KO β_1_-Halotag and MEF β_1_KO β_1_-Halotag paxillin-mCherry (both gifts from David Calderwood, Yale University). The former cell line was constructed by lentiviral transfection of a β_1_KO MEF cell line with an ecto-tag construct based on pLENTI expression vector and the latter as a consecutive lentiviral transfection with a paxillin-mCherry on pLENTI expression vector. The Halotag sequence was inserted on an exposed loop (β_1_ residues 91-114)^13^.

MEF β_1_KO β_1_-Halotag LifeAct-mScarlet that was constructed by lentiviral transfection of a β_1_KO β_1_-Halotag MEF cell line with a LifeAct-mScarlet on pLVX expression vector.

MEF β_1_KO β_1_-Halotag EMTB-iRFP that was constructed by lentiviral transfection of a β_1_KO β_1_-Halotag MEF cell line with an EMTB-iRFP on pLVX expression vector (gift from Simon De Beco, Paris Diderot University).

HeLa WT and HeLa β_1_-Halotag that we constructed by lentiviral transfection of a HeLa WT cell line with an ecto-tag construct based on pLENTI expression vector (gift from David Calderwood, Yale University).

MEF and HeLa cells were cultured in Dulbecco’s Modified Eagles Medium (DMEM) high glucose + GlutaMAX (Thermo Fischer Scientific) supplemented with 10% fetal bovine serum (FBS; EuroBio) and 1% penicillin-streptomycin (Thermo Fischer Scientific). Cells were cultured at 37°C in a humidified 5% CO_2_ atmosphere. Cells were routinely monitored for mycoplasma contamination following a well-established PCR-based method^14^ and found to be negative.

**DNA plasmids**

For transient cell transfections the following plasmids were used: talin-mCherry (Addgene plasmid #55137), vinculin-mCherry^15^, VASP-mCherry (Addgene plasmid #55151), kindlin-2-mCherry (gift from Christof Hauck, Konstanz University), zyxin-mCherry (gift from Danijela Vignjevic, Institut Curie).

For lentiviral transfections the following plasmids were used: β_1_-Halotag (gift from David Calderwood, Yale University), pLVX EMTB-iRFP (gift from Simon De Beco, Paris Diderot University) and pLVX LifeAct-mScarlet.

The pLVX LifeAct-mScarlet plasmid was prepared in two steps: a polymerase chain reaction using 5’- TCTAGAGCTACTAACTTCAGCCTGCTG-3’ / 5’- CGGTGGATCCCCTTCTTCC-3’ primers on Ibidi USA 60101 LifeAct-GFPtag2 plasmid was cloned with In-Fusion HD enzyme kit (Takara) into the pLVX vector (Clontech) digested with Not1 and BamH1 restriction enzymes (New England Biolabs). It was followed by a second In-Fusion HD cloning of a polymerase chain reaction using 5’-GAAGGGGATCCACCGATGGTGAGCAAGGGCGAGG-3’ / 5’- TTAGTAGCTCTAGACTTGTACAGCTCGTCCATGCC-3’ (mScarlet insert) using BamHI and XbaI (New England Biolabs).

**Cell transfections (transient by electroporation and stable lentiviral)**

Transient cell transfections were performed by electroporation following the previously described protocol^16^: trypsinized cells were resuspended at a concentration of 2.5 × 10^7^ cells/ml in 15 mM Hepes, pH 7.4, buffered medium. 200 μl of cell suspension was added to 50 μl of a solution containing 210 mM NaCl, 5 μg of plasmid DNA, and 30 μg of salmon sperm DNA carrier (Sigma Aldrich). Then cells were electroporated with a BioRad Gene Pulser at 950 μF and 240 V using 4-mm width cuvettes. Transiently transfected cells were analyzed after 48 h of the expression of the plasmid of interest.

Stable cell transfections were performed with lentiviral infections. Lentiviral particles (LVs) were produced in HEK 293T cells cultured in Dulbecco’s Modified Eagle’s Medium (Thermo Fisher Scientific), supplemented with 10% Fetal Bovine Serum (EuroBio) and 1% Pen/Strep + 1% Sodium Pyruvate + 1% Non-Essential Amino Acids solution (Gibco) at 37°C in 5% CO_2_ humidified incubators. Cells were plated the day before transfection in T75 flasks (approx. 6 million) to achieve 50–70% confluency the next day. Plasmids coding lentiviral components, pPAX2 (Gag-Pol-Hiv1) and pMDG2 (VSV-G), and the plasmid of interest at a ratio of 4:1:4 (µg), respectively, were transfected using PEI MAX 40k transfection reagent (Tebu-Bio) according to the manufacturer’s protocol. After 48 hrs, LVs were concentrated using a 100k Amicon column (Merck Millipore) and the pellet was resuspended up to 400 μL in PBS. MEF or HeLa cells were plated the day before infection in 6-well plates (approx. 100,000 cells) to achieve 50–70% confluency the next day. Cells were transduced with 100 μL of the desired LVs, for 72 hrs. After transduction, expressing cells were treated with 2 μg/mL puromycin. Positive cells were sorted using a SH800 FACS Cell Sorter (Sony).

**Integrin labeling and cell seeding in imaging chambers**

Cultured cells were serum starved for 24 hours before experiment. Then cells were detached by Versene solution (Sigma Aldrich) for 30 minutes at 37°C. Then they are incubated with an Alexa Fluor488 Halotag® ligand (Promega) (400 nM per 1 ml containing approximately 1.5-2 million cells) to label β_1_-integrin-Halotag for 10-15 minutes at room temperature. Then cells are span down at low centrifuge speed (1000 rpm in Megafuge 16R centrifuge from Thermo Scientific), resuspended in the cell buffer (25 mM HEPES pH 7.3; 120 mM NaCl; 7 mM KCl; 1.8 mM CaCl_2_; 0.8 mM MgCl_2_; 5 mM glucose) to remove the Versene and the excess of dye and filtered with Falcon 40 µm Cell Strainer (Corning) to remove cell clumps. Cells then were gently flown into the imaging chamber containing SLBs. The chamber is then sealed with mineral oil (M8410, Sigma Aldrich).

Since our aim was to study mechanosensitive aspects of cell adhesion, we avoided washing steps in chambers once cells were seeded. Indeed, flows in the chamber exert shearing forces on adhering cells that can reinforce cell adhesion^17^.

**Buffers, reagents, inhibitors**

The following cell buffer was used in imaging experiments: 25 mM HEPES pH 7.3; 120 mM NaCl; 7 mM KCl; 1.8 mM CaCl_2_; 0.8 mM MgCl_2_; 5 mM glucose.

SiR-tubulin (Cytoskeleton, Inc.) at the dilution suggested by the manufacturer was used to label microtubules in live cells.

In the experiments with manganese-treated cells, MnCl_2_ (Sigma Aldrich) was added (to the final 0.5 mM concentration) just before putting cells into the imaging chamber.

We used the following concentrations of inhibitor drugs in this study: 50 µM for the Arp2/3 complex inhibitor CK-666, 10 µM for the formin inhibitor SMIFH2 (Sigma Aldrich), 50 µM for the myosin inhibitor *p*-nitro-Blebbistatin (Cayman Chemical), 50 µM for the ROCK inhibitor Y27631 (Sigma Aldrich), 10 µM for Nocodazole (Sigma Aldrich) and 50 µM for the cytoplasmic dynein inhibitor Ciliobrevin D (Sigma Aldrich). For the live cell imaging experiments drugs were added at the same moment as cells were seeded to chambers. Control samples were treated with an equivalent amount of DMSO which did not exceed 0.001% v/v.

**Immunofluorescence**

For endogenous ELKS and KANK1 labeling, MEF cells were fixed in 100% ice-cold methanol (−20 °C, 5 min) followed by incubation in PBS with 1 mg ml^−1^ BSA (blocking buffer (BB)) all along the procedure. Fixed cells were washed in PBS and saturated in BB. Cells were incubated with the primary antibody (rabbit anti-ELKS^18^, 1:200, or rabbit anti-KANK1, Atlas antibodies, HPA005539, 1:200) diluted in BB (45 min), washed three times in BB, and incubated with the anti-mouse secondary antibody conjugated to Alexa Fluor 555 (Invitrogen) for 30 minutes. Cells were washed twice in BB, once in PBS and once in dH_2_O. Finally, coverslips were mounted in Abberior mounting medium (Abberior) and examined under fluorescence microscope.

**siRNA interference**

The sequences of siRNA for p150Glued are obtained from Eurogentec: GGUAUCUGACACGCUCCU and UAGGAGCGUGUCAGAUAC. Non-targeting siRNA ON-TARGETplus (D-001810-10-05) (Dharmacon, GE Healthcare) served as the siRNA control (“siRNA scramble”). siRNA transfection was performed using Lipofectamine RNAiMAX transfection reagent (Thermo Fischer) at a final concentration of 40nM for 48 hrs prior to cell imaging. Protein knock down efficiency was confirmed by real-time quantitative PCR and Western Blot analysis (Supplementary Fig. S6C).

**Western Blot analysis**

Western blots were performed on protein extracts of siRNA silenced HeLa to estimate the degree of knockdowns of Dynactin (p150glued) (Supplementary Fig. S6C). siRNA silenced Dynactin (p150glued) HeLa cells were resuspended in RIPA Buffer (50 mM TRIS pH8.0, 150 mM NaCl, 1% NP-40, 0,1% SDS) supplemented with cOmplete protease inhibitor cocktail (Roche). Sample was centrifuged at 20000 x g for 1 min and protein supernatant were collected and concentrations checked with Bradford assay. 100 µg of each sample was run on a 4-12% NuPAGE Gel, which was then transferred onto a PVDF membrane using the BioRad Trans-Blot Turbo system. Membranes were blocked in 5% Milk, TBST and treated with p150glued Antibody (Mouse, BD biosciences) or β-actin (Mouse, Genetex) as the loading control at a 1:1000 dilution in TBS-T at 4°C overnight, then the secondary Goat-anti-mouse HRP antibody (Sigma Aldrich) at 1:10 000 dilution in TBS-T, for 1h. Pierce™ ECL 2 Western Blotting Substrate (Thermo Scientific Kit) was added and blots developed using the Amersham Imager AI 680. To assess the degree of siRNA silencing of p150glued we quantified the relative change in the protein expression, using the Fiji “Measure” plugin and using the same region of interest (ROI) across the bands in one blot normalized by the intensity of the appropriate β–actin band (indicates the amount of loaded protein).

**Statistical analysis**

Statistical analysis was performed in GraphPad Prism. P<0.05 was considered statistically significant. At least three experiments were performed in order to conduct a statistical test (number of experiments are indicated as Nexp in the figure captions). Data sets were tested with the D’Agostino and Pearson normality tests. Normally distributed data sets were analyzed with Student’s t tests or with one-way ANOVA Tukey tests if they contained two or more conditions respectively. Data sets with non-normal distributions were analyzed with a Kruskal–Wallis test (multiple comparison) or Wilcoxon rank sum test (two-sample comparison).

**Bibliography (methods)**

1.	Nye, J. A. & Groves, J. T. Kinetic Control of Histidine-Tagged Protein Surface Density on Supported Lipid Bilayers. *Langmuir* **24**, 4145–4149 (2008).

2.	Humphries, J. D., Byron, A. & Humphries, M. J. Integrin ligands at a glance. *J. Cell Sci.* **119**, 3901–3903 (2006).

3.	Van Nhieu, G. T. & Isberg, R. R. The Yersinia pseudotuberculosis invasin protein and human fibronectin bind to mutually exclusive sites on the alpha 5 beta 1 integrin receptor. *J. Biol. Chem.* **266**, 24367–24375 (1991).

4.	Leong, J. M., Fournier, R. S. & Isberg, R. R. Identification of the integrin binding domain of the Yersinia pseudotuberculosis invasin protein. *EMBO J.* **9**, 1979–1989 (1990).

5.	Galush, W. J., Nye, J. A. & Groves, J. T. Quantitative fluorescence microscopy using supported lipid bilayer standards. *Biophys. J.* **95**, 2512–2519 (2008).

6.	Soumpasis, D. M. Theoretical analysis of fluorescence photobleaching recovery experiments. *Biophys. J.* **41**, 95–97 (1983).

7.	Guo, L. *et al.* Molecular Diffusion Measurement in Lipid Bilayers over Wide Concentration Ranges: A Comparative Study. *ChemPhysChem* **9**, 721–728 (2008).

8.	Pincet, F. *et al.* FRAP to Characterize Molecular Diffusion and Interaction in Various Membrane Environments. *PloS One* **11**, e0158457 (2016).

9.	Dahim, M. *et al.* Physical and photophysical characterization of a BODIPY phosphatidylcholine as a membrane probe. *Biophys. J.* **83**, 1511–1524 (2002).

10.	Nagle, J. F. & Tristram-Nagle, S. Structure of lipid bilayers. *Biochim. Biophys. Acta* **1469**, 159–195 (2000).

11.	Kapur, J. N., Sahoo, P. K. & Wong, A. K. C. A new method for gray-level picture thresholding using the entropy of the histogram. *Comput. Vis. Graph. Image Process.* **29**, 273–285 (1985).

12.	Sahoo, P., Wilkins, C. & Yeager, J. Threshold selection using Renyi’s entropy. *Pattern Recognit.* **30**, 71–84 (1997).

13.	Huet-Calderwood, C. *et al.* Novel ecto-tagged integrins reveal their trafficking in live cells. *Nat. Commun.* **8**, 570 (2017).

14.	Young, L., Sung, J., Stacey, G. & Masters, J. R. Detection of Mycoplasma in cell cultures. *Nat. Protoc.* **5**, 929–934 (2010).

15.	Valencia-Gallardo, C. *et al.* Shigella IpaA Binding to Talin Stimulates Filopodial Capture and Cell Adhesion. *Cell Rep.* **26**, 921-932.e6 (2019).

16.	Gautreau, A., Louvard, D. & Arpin, M. Morphogenic Effects of Ezrin Require a Phosphorylation-Induced Transition from Oligomers to Monomers at the Plasma Membrane. *J. Cell Biol.* **150**, 193–204 (2000).

17.	Riveline, D. *et al.* Focal contacts as mechanosensors: externally applied local mechanical force induces growth of focal contacts by an mDia1-dependent and ROCK-independent mechanism. *J. Cell Biol.* **153**, 1175–1186 (2001).

18.	Monier, S., Jollivet, F., Janoueix-Lerosey, I., Johannes, L. & Goud, B. Characterization of Novel Rab6-Interacting Proteins Involved in Endosome-to-TGN Transport. *Traffic* **3**, 289–297 (2002).
